# Generative Adversarial Networks Accurately Reconstruct Pan-Cancer Histology from Pathologic, Genomic, and Radiographic Latent Features

**DOI:** 10.1101/2024.03.22.586306

**Authors:** Frederick M. Howard, Hanna M. Hieromnimon, Siddhi Ramesh, James Dolezal, Sara Kochanny, Qianchen Zhang, Brad Feiger, Joseph Peterson, Cheng Fan, Charles M. Perou, Jasmine Vickery, Megan Sullivan, Kimberly Cole, Galina Khramtsova, Alexander T. Pearson

## Abstract

Artificial intelligence models have been increasingly used in the analysis of tumor histology to perform tasks ranging from routine classification to identification of novel molecular features. These approaches distill cancer histologic images into high-level features which are used in predictions, but understanding the biologic meaning of such features remains challenging. We present and validate a custom generative adversarial network – HistoXGAN – capable of reconstructing representative histology using feature vectors produced by common feature extractors. We evaluate HistoXGAN across 29 cancer subtypes and demonstrate that reconstructed images retain information regarding tumor grade, histologic subtype, and gene expression patterns. We leverage HistoXGAN to illustrate the underlying histologic features for deep learning models for actionable mutations, identify model reliance on histologic batch effect in predictions, and demonstrate accurate reconstruction of tumor histology from radiographic imaging for a ‘virtual biopsy’.

## Main

Histopathologic analysis of tumors is an essential step in the diagnosis and treatment of cancer in modern clinical oncology. The initial diagnosis of cancers depends on morphological assessment of biopsy samples and molecular profiling now informs prognosis and clinical therapeutic decisions in almost every cancer subtype. Machine learning, and more specifically deep learning, has been successfully applied to most standard steps of pathologic image analysis, and can segment^1^, diagnose^2^, grade^3^, and even predict recurrence or treatment response for tumors.^4^ As the field has evolved, studies have moved beyond basic pattern recognition towards identifying deeper disease traits and complex morphological features, including the identification of genomic and transcriptomic profiles directly from histology.^5–7^ Conceptually, deep learning models often condense complex visual information from histopathology into a small number of higher order features for prediction; often using pretraining from large image datasets like ImageNet^8^ or feature extractors trained with self-supervised learning (SSL)^9,10^. However, the opacity of these high level features limits adoption and deployment due to concerns about model trustworthiness^11^, and lack of interpretability limits the ability to gain new insight from the histologic patterns recognized by models.

A range of techniques exist for explaining machine learning model predictions in medical imaging, including saliency mapping, attention mechanisms, and perturbation-based approaches.^12,13^ However, these approaches may identify regions important for prediction such as tumor epithelium, but may not identify which characteristics of these regions have led to a positive or negative prediction.^13^ This is critically important in validation studies as results may be confounded by batch effects that are hard to characterize without thoroughly evaluating the features underpinning a model’s prediction.^14^ Generative adversarial networks (GANs) provide an attractive alternative framework for explainability. GAN frameworks like StyleGAN2 train a generator to produce realistic synthetic images able to fool a discriminator network and the resulting generator latent space captures semantic concepts and can be manipulated for intuitive image editing.^12^ Conditional GANs can be used to interpolate between two histologic classes – but training is time consuming and such an approach can only embed a limited number of classes.^5^

We consider an alternative approach to synthetic image generation - if histology could be reconstructed from high level features derived from SSL-based extractors, the visual meaning of these features (or models trained from these features) can be deciphered. Additionally, reconstruction of histology from base features enables development of accurate cross-modal autoencoders to reconstruct histology from other forms of data^15,16^, such as sequencing or MRI imaging, enabling a ‘virtual biopsy’. Approaches like Encoder4Editing and pix2style2pix allow the embedding of images into the latent space of a GAN but cannot successfully reconstruct histologic images in their base configurations.^17,18^ To address this, we present HistoXGAN, a modification of the StyleGAN2 architecture which uses a feature vector from highly effective SSL-based feature extractors as the basis of the GAN latent space to accurately reconstruct histology.

## Results

### Accurate Reconstruction of Histologic Structures

The HistoXGAN architecture is a modified StyleGAN2 generative adversarial network^12,19^, that ensures consistency of key image features extracted by SSL feature extractors during image generation^20,21^ (**Figure 1**, left; **Supplemental Figure 1**). In this way, generated images are structurally similar, but the location of image structures are allowed to vary between images. With this approach, images can be generated by providing a feature vector directly as the input latent vector, without requiring a separate encoder to project features into the StyleGAN latent space. We compared L1 loss / mean absolute error (**Figure 1**, middle) between extracted features from the input and reconstructed images generated by HistoXGAN and alternative encoders across 8,120 cases in the training TCGA dataset and an additional n = 1,328 cases from the Clinical Proteomics Tumor Analysis Consortium (CPTAC) dataset. HistoXGAN achieved the lowest reconstruction error, with a mean error of 0.034 (95% CI 0.034 – 0.034) across TCGA and 0.038 (95% CI 0.038 – 0.038) across CPTAC for reconstruction of CTransPath features (**Figure 2**, **Supplemental Table 1**), and a mean error of 0.010 (95% CI 0.010 – 0.010) across TCGA and 0.011 (95% CI 0.011 – 0.011) across CPTAC for reconstruction of RetCCL features (**Supplemental Figure 2**, **Supplemental Table 2**). This represented a 29% and 17% improvement over the second-best model, Encoder4Editing, for reconstruction of CTransPath and RetCCL features respectively. This improved feature reconstruction was reflected in pathologist review of images generated by HistoXGAN: eight tiles, one from each cancer subtype in the validation cohort, were regenerated using the four encoders, and four pathologists with over 50 years of combined experience were presented with the generated images in random order. The HistoXGAN reconstructions were judged as most similar to input images in 75% (24 / 32) of cases when using CTransPath features and 84% (27 / 32) of cases using RetCCL features (**Figure 2B**, **Supplemental Figure 2B**).

**Figure 1.**
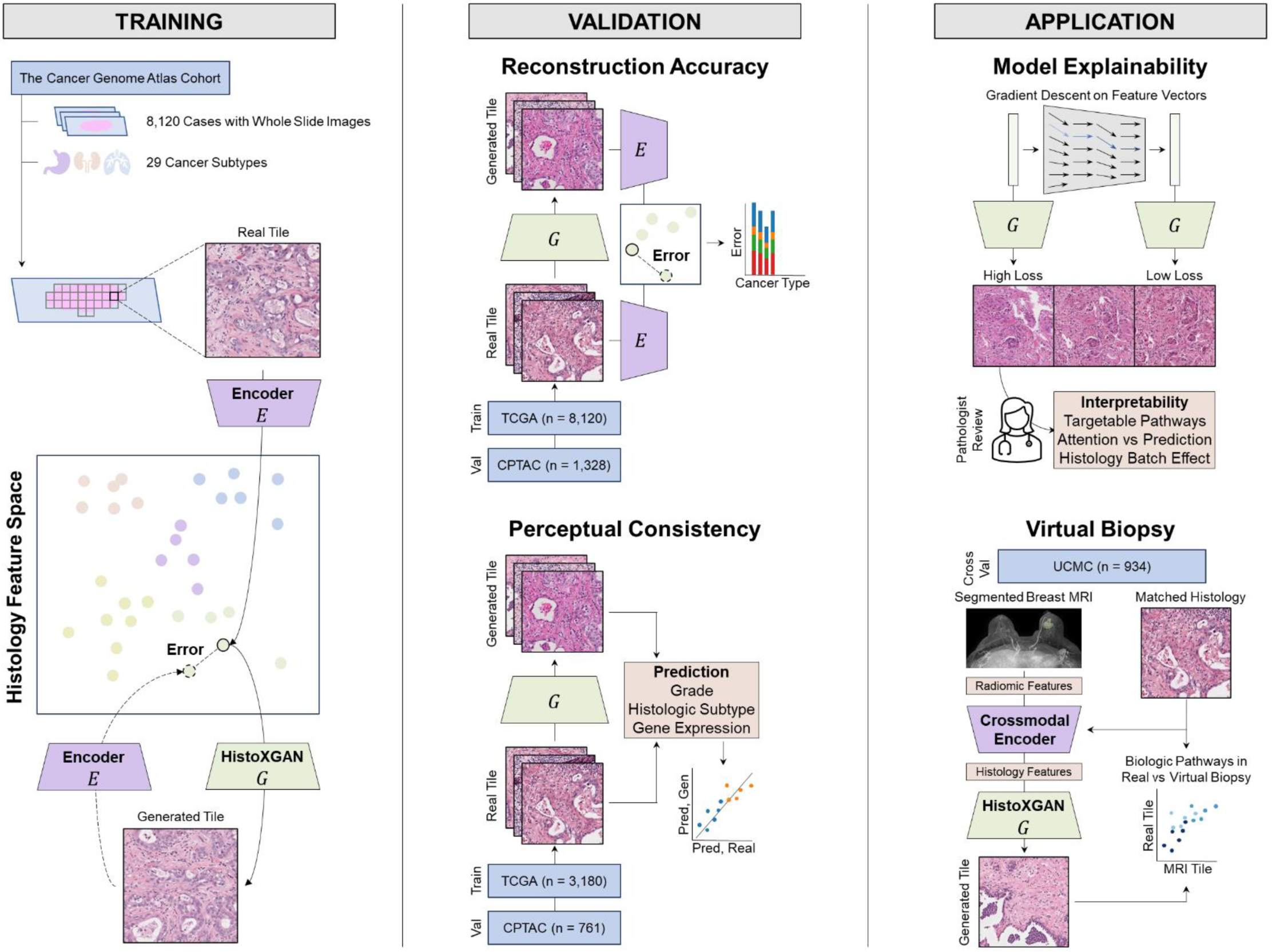
Overview of HistoXGAN Training, Validation and Application. **Left.** The HistoXGAN generator *G* was trained using 8,120 cases across 29 cancer types in The Cancer Genome Atlas (TCGA). HistoXGAN takes as an input a histologic feature vector derived from any self-supervised feature extractor *E*, and generates a histology tile with near-identical features with respect to the same feature extractor. **Middle.** In this study, we demonstrate that this architecture accurately reconstructs the encoded features from multiple feature extractors in both the training TCGA dataset and an external dataset of 3,079 slides from 1,328 cases from Clinical Proteomic Tumor Analysis Consortium (CPTAC). Additionally, we demonstrate the real and reconstructed images carry near-identical information of interpretable pathologic features such as grade, histologic subtype, and gene expression data. **Right.** We showcase the applications of this architecture for model interpretability, using gradient descent to illustrate features used in deep learning model predictions. Through systematic review of these features with expert pathologists we identify characteristics of cancers with targetable pathways such as homologous recombination deficiency and PIK3CA in breast cancer, and illustrate application for attention based models. Finally, we train a crossmodal encoder to translate MRI radiomic features into histology features using paired breast MRIs and histology from 934 breast cancer cases from the University of Chicago Medical Center, and apply HistoXGAN to generate representative histology directly from MRI.

**Figure 2.**
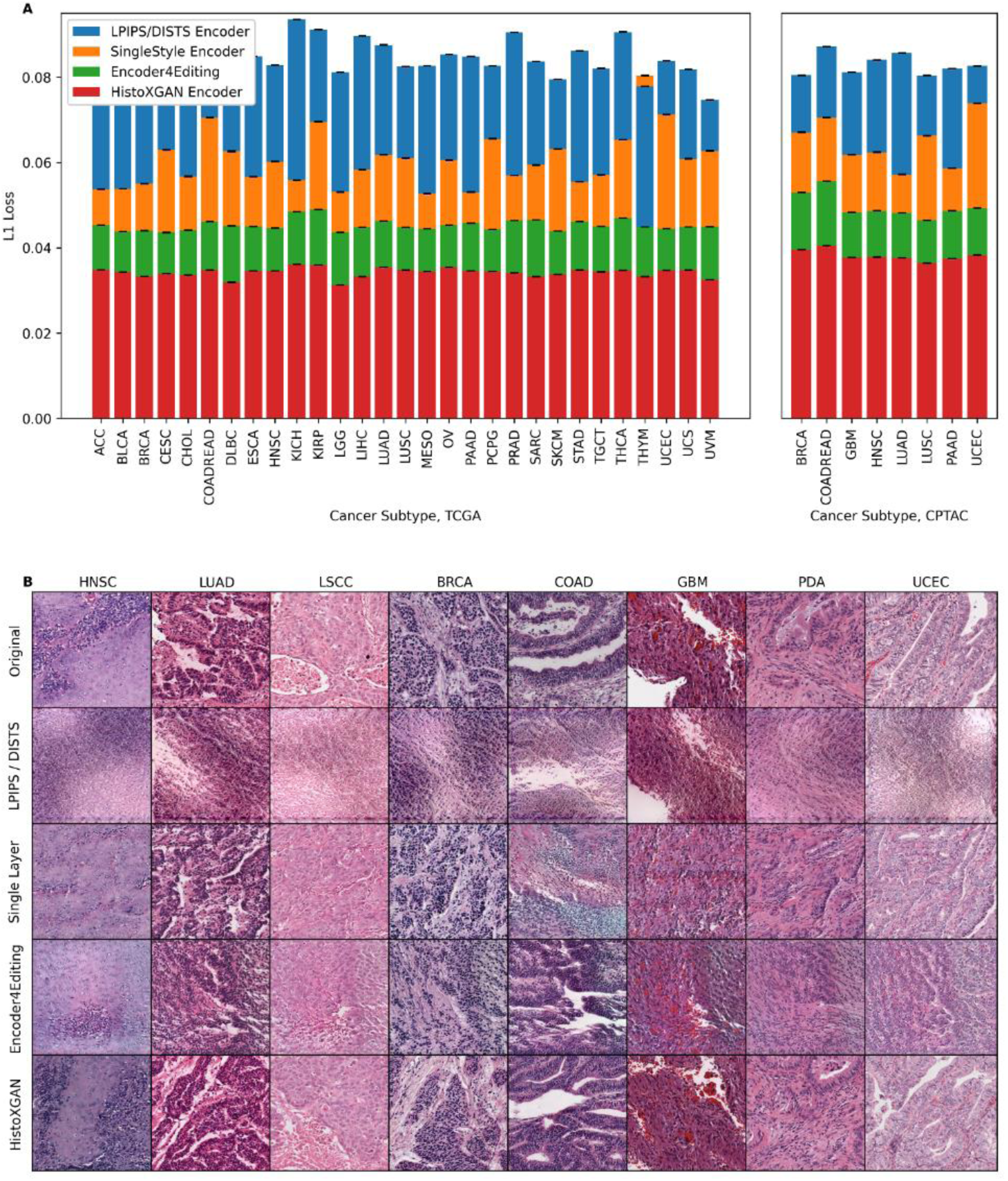
Reconstruction Accuracy in Training and Validation Datasets for CTransPath Encoders. We compare reconstruction accuracy from the real and reconstructed images for HistoXGAN and other architectures for embedding images in GAN latent space. For comparison, we use encoders designed to recreate images from a StyleGAN2 model trained identically to the HistoXGAN model. The Learned Perceptual Image Patch Similarity (LPIPS) / Deep Image Structure and Texture Similarity (DISTS) encoder uses an equal ratio of LPIPS / DISTS loss between the real and reconstructed images to train the encoder. The Single Layer and Encoder4Editing encoders are trained to minimize L1 loss between CTransPath feature vector of the real and reconstructed images. **A.** HistoXGAN provides more accurate reconstruction of CTransPath features across the TCGA dataset used for GAN training (n = 8,120) and CPTAC validation (n = 1,328) dataset, achieving an average of 30% improvement in L1 loss over the Encoder4Editing encodings in the validation dataset. **B.** HistoXGAN reconstructed images consistently provided more accurate representations of features from the input image across cancer types in the CPTAC validation dataset.

### Reconstructed Histology Retains Meaningful Representations of Tumor Biology

A meaningful synthetic reconstruction of tumor histology from features in a shared latent space should retain key elements of tumor biology that are reflected in pathology – for example, the tumor grade of the reconstructed histology should be identical to the original. To test these aspects of our approach to reconstruction in a systematic and quantitative fashion, we trained deep learning models for grade (TCGA/CPTAC n = 943/100 breast, 168/139 pancreas, 391/107 head and neck, and 227/none prostate), histologic subtype (743/92 breast, 941/415 lung, 147/none esophageal, and 363/none kidney), and gene expression (941/97 breast), and compared predictions made from whole slide images to those made from the same set of tiles reconstructed with HistoXGAN (**Figure 1B**, middle). Importantly, models for these tasks were trained with a non-SSL based architecture, so predictions are not based on the same image features used to train HistoXGAN. For prediction of high tumor grade (defined as grade 3 for breast / pancreatic, grade 3 or 4 for head and neck, or Gleason grade 9 or 10 for prostate; **Figure 2**, **Supplemental Table 3**), predictions from real slides and from reconstructed tiles were highly correlated with correlation coefficients from ranging from 0.52 (95% CI 0.36 – 0.65, p = 2.33 × 10^−8^) in CPTAC-BRCA to 0.85 (95% CI 0.79 – 0.89, p = 1.01 × 10 ^−29^) in CPTAC-HNSC. Similar findings were seen for the prediction of tumor histologic subtype in TCGA/CPTAC breast, lung, esophageal, and renal cancers (**Supplemental Figure 3**, **Supplemental Table 4**), as well as for prediction of gene expression of *CD3G*, *COL1A1*, *MKI67*, and *EPCAM* in TCGA/CPTAC breast cancer cohorts (**Figure 2**, **Supplemental Table 5**). Transition between states of grade, histology, and gene expression can all be readily visualized using HistoXGAN reconstructions (**Figure 2A and C**, **Supplemental Figure 3A**).

### Interpretability of Models and Applications to Understand Tumor Biology

We demonstrate the utility of HistoXGAN to leverage deep learning models to uncover new insights by characterizing histologic manifestations of tumor biology. Given the increasing number of targetable molecular alterations predicable from histology^9^, we evaluated the explainability of pathways with targeted therapies in breast cancer that have not been thoroughly explored – namely, *PIK3CA* alterations and homologous recombination deficiency (HRD). Models were trained to predict these alterations in TCGA-BRCA, achieving an average AUROC of 0.61 (n = 901, range 0.58 – 0.63) for *PIK3CA* alteration and 0.71 (n = 820, range 0.65 – 0.76) for high HRD score respectively on three-fold cross validation, similar to previously published models. Image tiles were altered through gradient descent (**Figure 1**, right) to maximize the predictions for *PIK3CA* mutation and HRD (**Figure 4 A & B**), and a set of 22 transitions was reviewed for qualitative analysis of nuclear, cytoplasmic, stromal, immune, and vascular features by four pathologists specializing in breast pathology. Transition to *PIK3CA* mutation (**Figure 4C**) was morphologically associated with increase in abundance (in 45% of transitions) and eosinophilic appearance of cytoplasm (in 68%), increased tubule formation (in 36%), increased invasion into stroma (in 63%) and decreased nuclear to cytoplasmic ratio (in 18%) with variable changes in nuclear size. Transition to high HRD score (**Figure 4D**) was associated with prominent nucleoli (in 59%), nuclear crowding / increased nuclear density (45%) with larger (36%) pleomorphic nuclei (18%) with occasional multinucleated cells (9%), increased lymphocytosis (54%) and tumor cell necrosis (5%).

**Figure 3:**
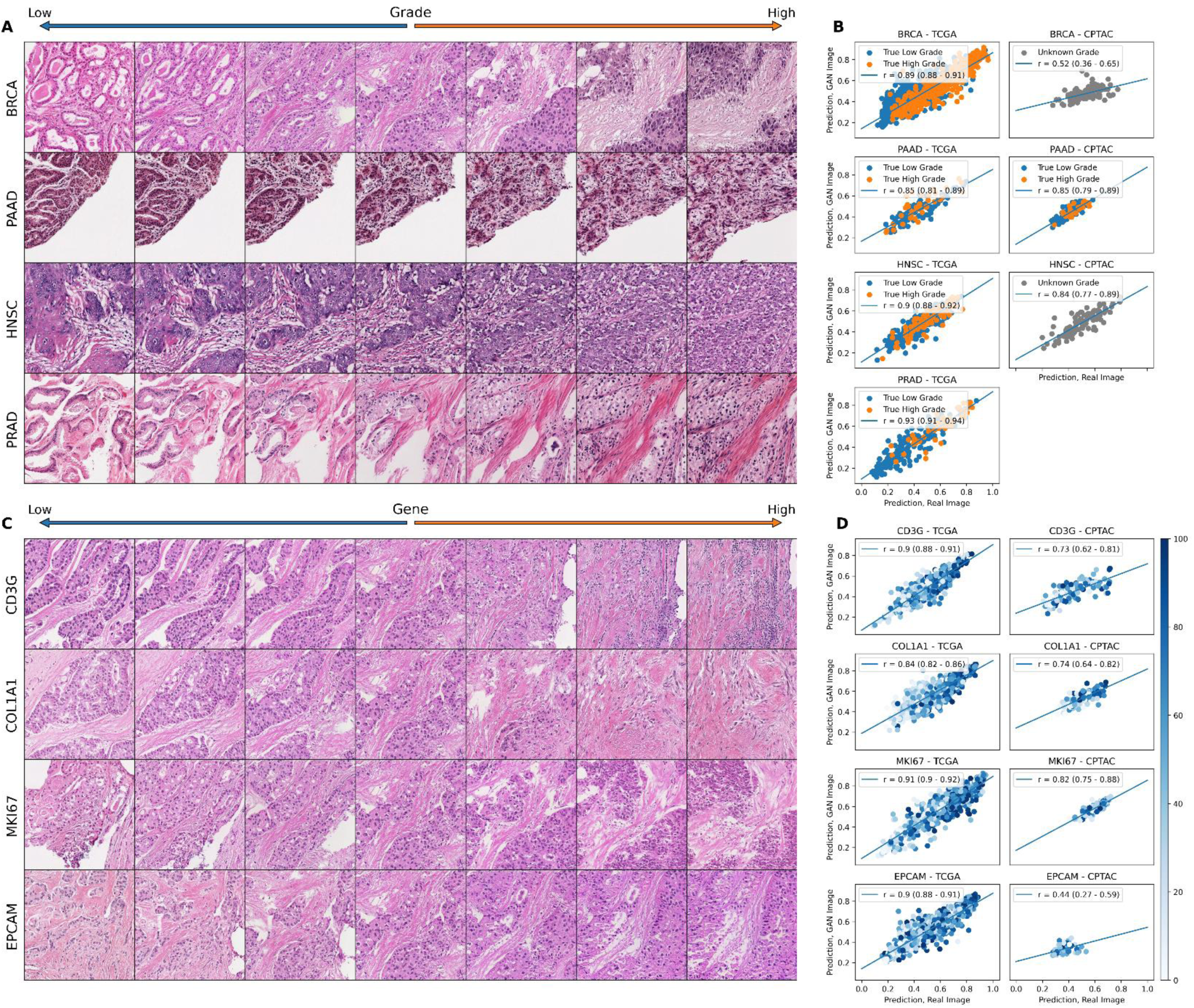
Perceptual Consistency of Tumor Grade and Gene Expression in Reconstructed Images. **A.** Illustration of transition between low and high grade (defined as grade 3 for breast and pancreas, grade 3 or 4 for head and neck, and Gleason grades 9 or 10 for prostate) across a single image from four cancer types. A vector representing high grade is derived from the coefficients of a logistic regression predicting grade from CTransPath features. This vector is subtracted from the base image to visualize lower grade, and added to the base image to visualize higher grade. **B.** Correlation between predictions of grade from real and reconstructed tiles, averaged per patient, across cancer types, demonstrating a high perceptual similarity of the grade of the real and generated images. For the TCGA datasets, a deep learning model was trained to predict grade from real tiles for each cancer type using three-fold cross validation. The correlation between predictions for real / generated images is aggregated for the three held out validation sets. For the CPTAC validation, a deep learning model trained across the entire corresponding TCGA dataset was used to generate predictions. True pathologist determined high versus low grade is annotated on the images when available. **C.** Illustration of transition between expression of select genes across a single image from TCGA-BRCA. **D.** Correlation between predictions of gene expression from real and reconstructed tiles, averaged per patient, demonstrating a high perceptual similarity of the gene expression of the real and generated images. True gene expression as a percentile from 0 – 100 is indicated by the color of each data point.

**Figure 4.**
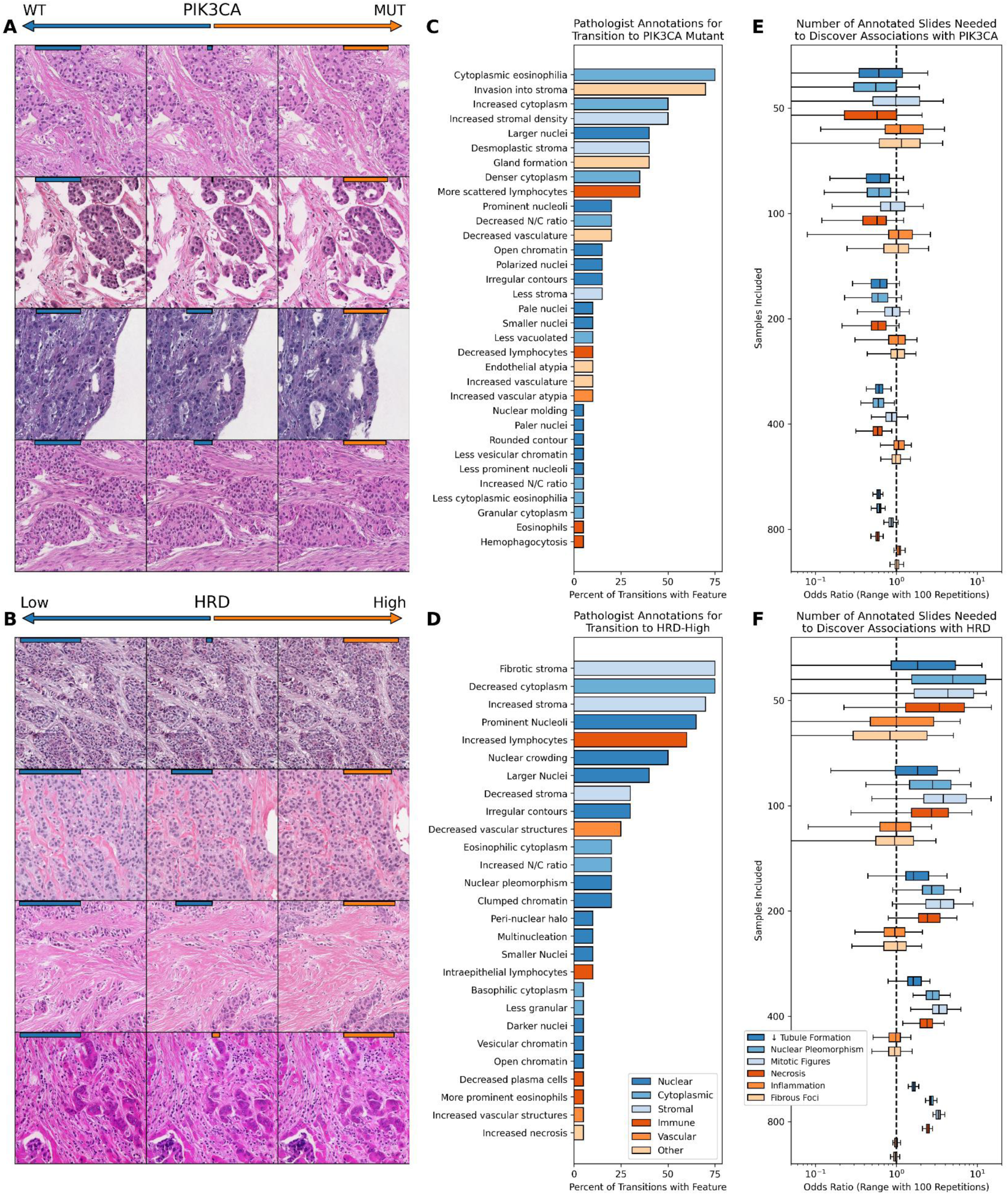
Illustration of Model Predictions for Targetable Alterations in Breast Cancer. Tile based weakly supervised models were trained to predict *PIK3CA* mutations and homologous recombination deficiency (HRD) across the entire TCGA-BRCA dataset (n = 963 for *PIK3CA*, n = 871 for HRD), as these two pathways are common and have FDA approved therapies available. Feature vectors were extracted along random baseline image tiles, and gradient descent was used to adjust the feature vector to be more / less likely to be predicted as *PIK3CA* altered / homologous recombination deficient. This transition was repeated for 20 images, which were reviewed by four pathologists for qualitative assessment. Strength of the base model prediction for each outcome (*PIK3CA* altered versus wild-type and homologous recombination deficient versus proficient) is illustrated with orange / blue bars on top of each image, confirming that gradient descent successfully transitions between model predictions for each base image. **A.** Transition to *PIK3CA* alteration were morphologically associated with increase in abundance and eosinophilic appearance of cytoplasm and stroma, increased tubule formation, and decreased nuclear to cytoplasmic ratio with variable changes in nuclear pleomorphism. **B.** Transition to high HRD score was associated with nuclear crowding and pleomorphism with occasional multinucleated cells, an increased nuclear / cytoplasmic ratio, increased lymphocytosis, and tumor cell necrosis. **C & D.** Structured pathologist review of 20 transitions from low to high model prediction highlight features that were associated with the selected genomic alterations. **E & F.** To determine how many histology slides in TCGA would be needed to be fully annotated through traditional methods to uncover some of these same associations, adjusted odds ratio of the association of histologic features with *PIK3CA* and HRD status are plotted as a function of available slides. Odds ratios were estimated through 100 iterations of random sampling of the listed number of slides, with box plots indicating the distribution. Histologic features such as increased tubule formation in *PIK3CA* and necrotic tumor in homologous recombination deficiency which were evident on review of 20 image transitions would require annotation of up to 400 slides with traditional histologic review to uncover statistically significant associations.

To validate these findings, we compared annotations for epithelial, nuclear, and mitotic grade as well as annotations for necrosis, lymphocytosis, and fibrous foci across TCGA-BRCA between *PIK3CA* mutant and wild type, and HRD-high and low tumors, generally yielding consistent findings (**Supplemental Tables 6 and 7**). To illustrate the benefit of synthetic histology to power discovery, we determined the number of samples required to identify significant associations of mutational status with these annotations. Some pathologic features, such as increased tubule formation in *PIK3CA* mutant tumors, or immune infiltrate in HRD-high tumors would require over 200 whole slide images annotated to demonstrate a significant association, whereas pathologic review of just 22 image transitions clearly uncovered these associations in this study (**Figure 4 E & F**).

Furthermore, the emergence of attention-based multiple-instance learning (MIL) necessitates approaches that can disentangle features used for attention vs outcome prediction to truly facilitate interpretability. We demonstrate an application of HistoXGAN to attention-MIL models – using gradient descent, feature vectors can be perturbed to increase or decrease model attention and model prediction separately, allowing independent visualization along these two axes (**Supplemental Figure 4**). Applying this approach to models trained to predict grade and cancer subtype illustrates that low attention for these models is associated with benign appearing fibrous tissue.

### Applying Generative Histology to Enable a Virtual Tumor Biopsy

Radiomic analysis of MRI images have been applied to predict key histologic features such as tumor grade and immune infiltrates. However, by predicting histologic SSL feature vectors from radiomic features, a representative tumor image can be reconstructed for downstream analysis, representing a ‘virtual tumor biopsy’. With five-fold cross validation across 934 cases with paired MRI and histology, we trained encoders to predict the SSL pathology feature vectors from radiomic features using and pooled the predicted features from the held-out test sets for analysis (**Figure 5**). Across these test sets, a mean L1 error of 0.078 (95% CI 0.077 – 0.079) in reconstruction of the histologic feature vectors was observed, comparable to the mean L1 error between pairs of tile images within the tumor across slides (0.074, 95% CI 0.073 – 0.075), and lower than the average inter-patient difference between average feature vectors of 0.092 (95% CI 0.092 – 0.093, **Figure 5B**). To understand how accurately the recreated images represent true histology across a wide range of meaningful biologic features, we used models pretrained in TCGA to predict grade, histologic subtype, and 775 putatively important gene expression signatures in breast cancer which can be accurately predicted from histology (average Pearson r between true gene signature and histology prediction of 0.45, range 0.09 – 0.74; average FDR-corrected p-value for correlation of 1.10 × 10^−5^; **Supplemental Figure 5**, **Supplemental Table 8**). Of note, predictions from these models largely fell into a smaller number of orthogonal categories (**Figure 5C**). We found significant correlation between predictions from real histology slide and predictions from virtual biopsy tiles for 213 of these 777 features after FDR correction. Accurate predictions were seen for IFNγ3 (Pearson r 0.21, 95% CI 0.15 – 0.28, corrected p = 4.5 × 10^−11^) as well as multiple breast cancer prognostic signatures^22,23^, including a research-based version of OncotypeDX recurrence score (r 0.15, 95% CI 0.08 – 0.21, p = 6.9 × 10^−6^, **Supplemental Table 9**). Although the correlation coefficients for these predictions are not high, the number of positively correlated signatures suggests some elements of true tumor biology are present in these virtual biopsies. Additionally, we found that accurate prediction was largely limited by the accuracy of the radiomic features themselves - a logistic regression trained to predict the result of these 777 features directly from radiomic features performed similarly to reconstructed histology (**Figure 5D**).

**Figure 5.**
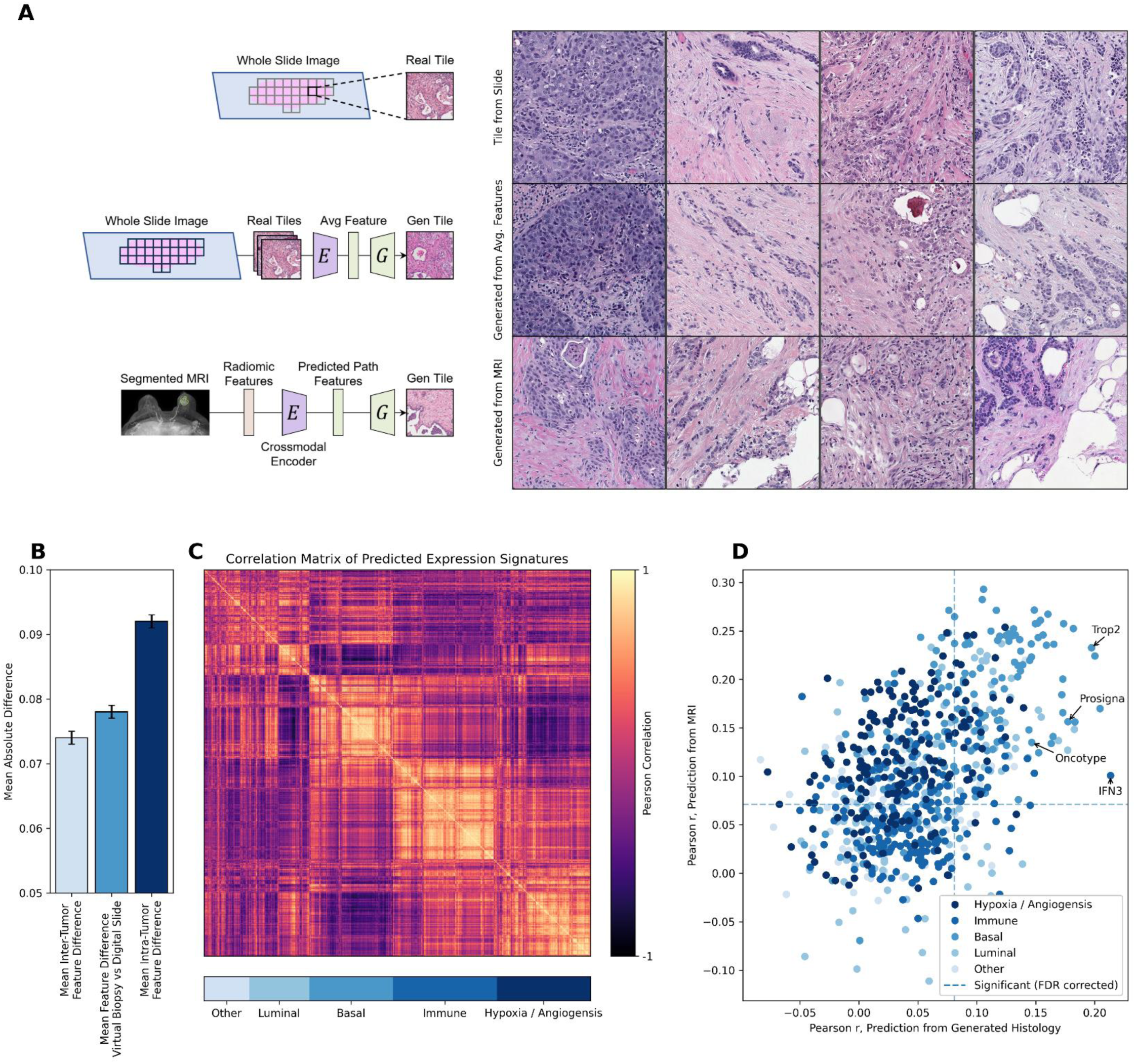
A ‘Virtual Biopsy’ Reconstructing Histology from Radiomic Features. An encoder was trained to predict the average SSL histology feature vector, using 16,379 radiomic features extracted systematically from 934 breast tumors with paired MRI and digital histology available. Five-fold cross validation was performed, with predictions pooled from the held out test set. **A.** Representative images from the scanned histology slide, reconstruction of the image from an average SSL histology feature vector, and reconstruction directly from radiomic features (predictions made using cases from the test set for each fold). **B.** To illustrate the accuracy of feature reconstruction from MRI, we compare the average difference of features extracted from these virtual biopsy images from the corresponding average tumor features from whole slide images (middle column), to the mean difference between image tiles within tumors (left), and the mean difference between image tiles between tumors (right). Although the generated virtual biopsy image is not indistinguishable from other images within the same tumor, the error is much lower than image tiles selected from other tumors, suggesting homology between the virtual biopsy and original tumor. **C.** To explore the ‘biologic accuracy’ of generated images, we used models trained in TCGA to predict 775 RNA sequencing features, as well as pathologist annotations of grade and histologic subtype. Predictions from these models largely fell into five orthogonal categories, with correlation between model predictions across the true 934 breast tumors in the dataset shown. **D.** We present the correlation between predictions for these 777 features from the true whole slide images versus pathology tiles generated from MRI features. As a comparator, we evaluated the accuracy of a logistic regression model to predict these features directly from MRI (without the intermediary of generated histology). The accuracy of prediction of features from generated histology was largely dependent on accuracy of prediction from MRI (i.e., features could not be predicted from histology if they could not be predicted from MRI). However, a number of features from the luminal / basal axis, including tumor grade, could be predicted.

### Identifying Models Influenced by Histologic Batch Effect

Some features predictable from histology using deep learning can be attributable to batch effect, or non-biologic differences that arise due to slide staining, tissue processing, image resolution, or other differences between batches of cases from model training and evaluation. To deconvolute the features used in model prediction, we can apply principal component analysis to the gradients generated from deep learning models for a large set of input images, to generate orthogonal feature vectors representing the directions traveled to increase / decrease model prediction. We first apply this approach to models trained to predict grade (true biologic feature) and contributing site (attributable entirely to batch effect) in TCGA-BRCA (n = 934). Principal components are sorted by the relative contribution to the difference in gradients towards an increased / decreased prediction. Whereas a number of distinct components comprise the prediction of higher grade, 69% of the difference in gradient for site prediction is comprised of a single component representing a change in tissue stain pattern, whereas the highest contribution from a single component for grade is 20% (**Supplemental Figure 6**, **Supplemental Table 10**). We demonstrate this effect is consistent for site prediction across tumor types with an average of 54% of site prediction attributable to a single component with a strong color variation (**Figure 6A**), with a similar visual pattern seen for ancestry prediction (**Figure 6B**). Reinhard normalization does not eliminate the dependence on a single color pattern with 37% of predictions remaining attributable to a single component – although the color pattern of prediction is inverted, likely due to the overcorrection / introducing color changes of image background elements (**Figure 6C**). CycleGAN normalization results in an improvement in the dependence of prediction on stain color (**Figure 6D**).

**Figure 6.**
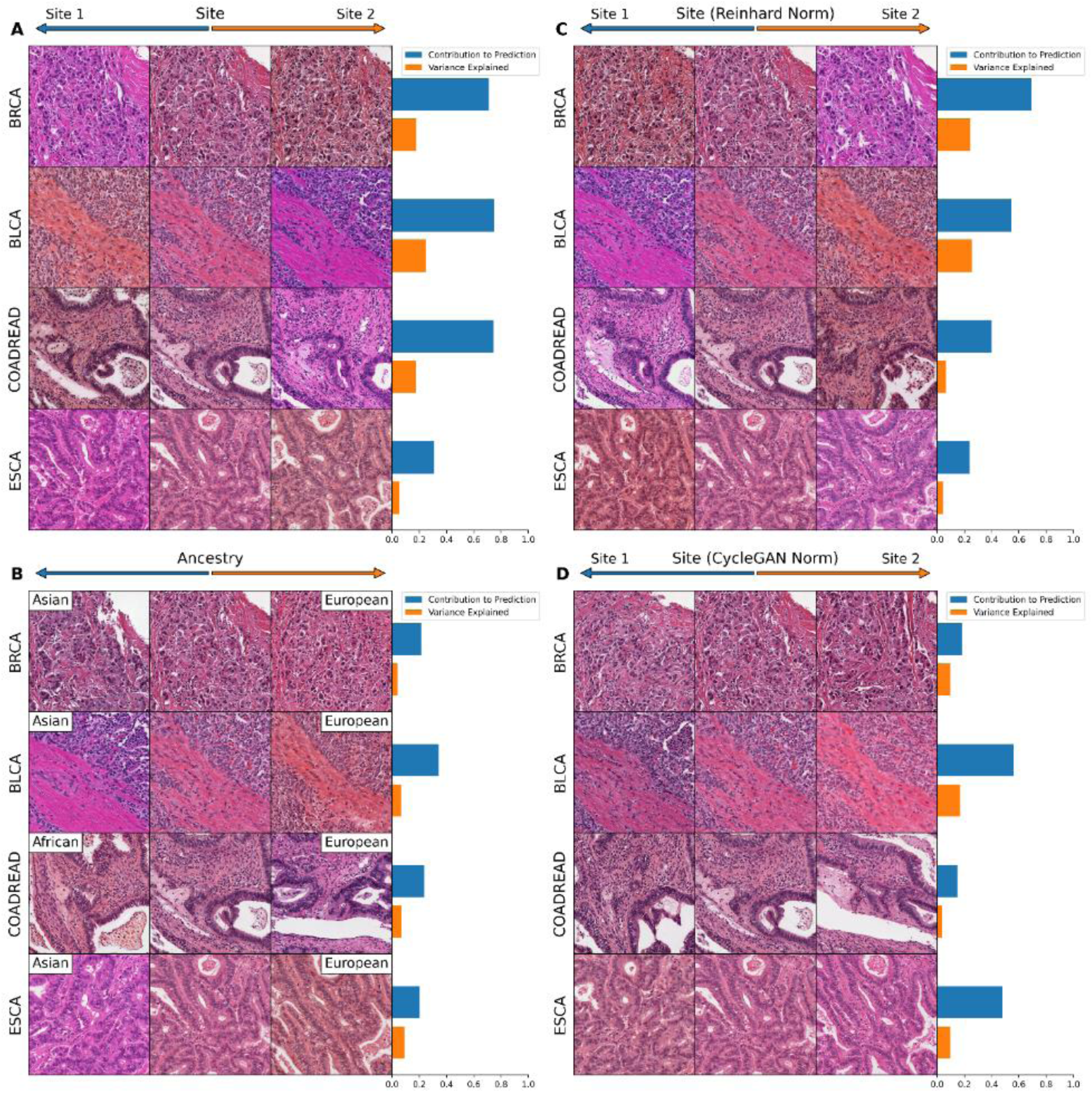
Visualizing Histology Batch Effect and Mitigation with Normalization. Tile based weakly supervised models were trained to predict tissue source site and patient ancestry class (a batch confounded outcome) across select cancer subtypes in TCGA. The gradient with respect to a prediction of these outcomes was calculated for the average feature vector across each slide in the dataset. Principal component analysis was applied to these gradients, and components were sorted by the magnitude of difference of the component between gradients toward each outcome class. Results are then illustrated for this first principal component (i.e. the component contributing most to model prediction). **A.** Model predictions for source site are highly homogenous, with an average 54% of the difference in gradients due to the first principal component. Perturbation of images along this component illustrate that it largely represents change in the staining pattern of slides. **B.** Slide stain patterns also contribute to prediction of ancestry, although this first principal component constitutes a smaller proportion of gradients. **C.** Reinhard normalization does not eliminate the impact of stain pattern on prediction of site, although it leads to an inversion of the stain detected by the model, perhaps due to overcorrection during normalization. **D.** CycleGAN normalization reduces the dependence of predictions on a single principal component, and this most predictive component is no longer clearly indicative of staining differences.

## Discussion

This study presents HistoXGAN, a novel generative adversarial network architecture for digital pathology image reconstruction that preserves interpretable disease traits. By integrating recent SSL pathology feature extractors^20,21^ with a modified StyleGAN2 generator, HistoXGAN allows interactive manipulation of tissue morphology while maintaining crucial architectural elements indicative of underlying tumor biology. Quantitative analysis across over 11,000 images validates that HistoXGAN reconstructions accurately recapitulate pathological grade, subtype, and gene expression patterns. Expert pathologist review further corroborates that generated images exhibit superior perceptual similarity to the original histology compared to other modern encoders^24,25^.

In recent years, deep learning has been applied to predict molecular features of cancers directly from histology with varying degrees of accuracy^6,9,26^ – but understanding the basis of these predictions remains challenging. Conditional GANs have been used to understand clear-cut histologic features, but must be retrained for each class comparison and cannot demonstrate multiple transitions simultaneously^27^. As HistoXGAN accurately recapitulates histologic features that are easily interpreted by pathologists like cancer grade and subtype, it can likely be applied for discovery of histologic patterns associated with molecular pathways. We applied HistoXGAN to prediction of predictable, actionable alterations in breast cancer – including *PIK3CA* mutation^28,29^ and HRD status (as defined by high HRD score)^30,31^. Prior studies evaluating histologic features of *PIK3CA* mutations have described conflicting findings – with one study reporting low grade^32^ and others describing sarcomatoid features with areas of high grade carcinoma^33^. We demonstrate that deep learning prediction of *PIK3CA* was associated with more well differentiated tubule formation and increased cytoplasm to nuclear ratio, but also increased nuclear size / pleomorphism which may explain the conflicting findings regarding grade in prior studies. Similarly, HRD status and BRCA alteration has been associated with high tumor cell density, with a high nucleus/cytoplasm ratio and conspicuous nucleoli, laminated fibrosis, and high TIL content as well as regions of hemorrhagic suffusion associated with necrotic tissue.^34,35^ Review of HistoXGAN images by specialized breast pathologists revealed associated visual features, including nuclear crowding and pleomorphism, an increased nuclear to cytoplasmic ratio, tumor-infiltrating lymphocytes, and areas of tumor necrosis, all of which are consistent with the aforementioned published results. Overall, these findings add confidence that there are true biologic features identified by deep learning models for these alterations, and these features can be used to identify patients at higher likelihood of molecular alterations in standard histologic analysis of tumors. With the rapid growth studies using AI for pathologic image analysis, HistoXGAN may be an important tool to ensure models predictions are based on rational biologically relevant features.

Additionally, the emergence of attention-based multiple-instance learning necessitates approaches that can disentangle features used for attention versus outcome prediction to facilitate true model interpretability. In general, publications have presented heatmaps of model attention, or selected high/low prediction tiles to illustrate potential features used in prediction^10,36^. We demonstrate an elegant application of HistoXGAN to attention-MIL models - using gradient descent, feature vectors can be perturbed to increase or decrease model attention and predictions separately. This enables independent visualization along these two axes, which is critical for understanding if predictions are driven by meaningful morphology versus dataset biases that attract model attention. Applying this methodology to grade and subtype prediction models illustrates that low model attention correlates with benign fibrous tissue, rather than malignant elements, verifying attention is applied to tumor regions. Overall, this approach can discern if attention mechanisms highlight biologically salient regions, or if predictions are partially confounded by irrelevant features that draw attention. Furthermore, we demonstrate that models that are highly confounded by site-specific factors such as ancestry can be quickly identified with HistoXGAN^14,37^. Interestingly, standard stain normalization such as the Reinhard^38^ method demonstrate ‘overcorrection’ of color, as HistoXGAN illustrates models trained after Reinhard normalization identify the inverse color transition as associated with site, whereas CycleGAN normalization^39^ was much more effective at eliminating learned staining patterns of tissue submitting sites.

Furthermore, studies have applied cross-modal autoencoders to understand common ‘latent spaces’ between multiple forms of data, but these approaches have not been performed with digital histology^15,16^. In particular, models to predict the histologic diagnosis or cancer phenotypes from imaging have been described as ‘virtual biopsies’^40–43^ without the intermediate step of tissue histology reconstruction. We demonstrate here that HistoXGAN can be used to create realistic representations of tumor histology directly from imaging radiomic features, and that biologic elements of tumor aggressiveness can be identified from the recreated pathology images. This approach could also be used for explainability of radiomic predictions – for example, if a radiomic feature is predictive of response to therapy, histology could be generated as a function of this feature to determine if the radiomic feature is highly correlated with known pathologic predictors of response such as tumor grade. As opposed to prior virtual biopsy approaches, generating a representative section of H&E histology theoretically allows for characterization of any pathologic feature that could be performed from a true biopsy, rather than restricting analysis to a limited set of outcomes used during training. With a larger training dataset across multiple cancer types, this tool could theoretically be applied as a first step to cancer lesion diagnosis in areas that are inaccessible, or as a quality control check, where if biopsy results are discordant with predicted histology it may suggest inadequate sampling and need for re-biopsy.

Several opportunities exist to build upon this study’s limitations. HistoXGAN was developed using histology across the TCGA dataset with predominantly solid tumors, and reconstructive accuracy may be limited for cancers that are not well represented within this dataset – such as hematologic malignancies or rare tumor subtypes. Similarly, this approach may not reconstruct representative histology for benign neoplasms or lesions not represented in TCGA. Our ‘virtual biopsy’ approach focused on characterizing cases with known breast tumors, and similar encoders would need to be trained with cases of benign disease if such a tool was used to distinguish malignant potential of lesions. Using generative approaches for discovery of biologic pathways requires a high degree of confidence in the accuracy of the generative model; for example, the identified pathologic characteristics associated with *PIK3CA* alteration and HRD status are relatively subtle, and it can be difficult to verify the accuracy of features derived from generative images although they are consistent with prior reports. However, the fact that known pathologic features such as grade and histologic subtype are accurately encoded by HistoXGAN provides some confidence that this approach can be used to describe histologic associations with rare mutations that have not yet been fully characterized.

In summary, this work presents HistoXGAN, an architecture integrating self-supervised learning and generative adversarial networks to facilitate interpretable manipulation of digital pathology images while maintaining important disease-specific morphological traits. Evaluations in over 11,000 images demonstrate quantitatively accurate reconstructions as well as qualitative expert pathologist endorsements of similarity. This technology can greatly aid in the interpretability of artificial intelligence models, discover novel biologic insights into targetable pathways to accelerate biomarker development, and even be leveraged to non-invasively sample cancer histology for a true ‘virtual biopsy’.

## Methods

### Data Sources and Image Extraction

Patient data and whole-slide images were selected from 29 tumor type datasets from TCGA (n = 8,120) and eight corresponding tumor types from CPTAC (n = 1,890) were used for model validation. Slides and associated clinical data were accessed through the Genomic Data Commons Portal (https://portal.gdc.cancer.gov/). Ancestry was determined using genomic ancestry calls from the work published by Carrot-Zhang and colleagues^44^. Annotations for HRD score were taken from Knijnenburg et al, and binarized at a score of ≥ 42 for training of HRD models^45^. The cohort of 934 matched pairs of tumor histology / MRI images was collected from University of Chicago under IRB approved protocol 22-0707 from patients diagnosed from 2006 – 2021 (**Supplemental Table 11**). Slide images were extracted using the Slideflow pipeline with a tile size of 512px per 400 µm and filtering to remove tiles with > 60% grayspace^46^. For GAN training and for applications with weakly supervised models without an attention component, slides were only extracted within pathologist annotated tumor regions of interest. For attention-based multiple instance learning (MIL) models, tiles were extracted from unannotated slides.

### Generative Adversarial Network and Encoder Training

The HistoXGAN architecture is a custom version of StyleGAN2, comprised of a generator *G*, discriminator *D*, and an encoder *E* (**Supplemental Figure 1**). Model training consists of two important modifications to the StyleGAN2 architecture. First, the latent vector *z* used for each batch during the generator training is replaced with the feature vector extracted by the SSL encoder. Second, a weighted L1 loss comparing the SSL extracted image features from the real image to those extracted from the generated image is added to the generator loss.

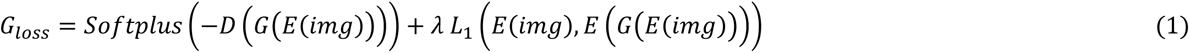

The HistoXGAN and StyleGAN2 networks used in this study were trained with 25,000,000 images across the entire TCGA dataset with a batch size of 256, with a lambda weight of 100 for the above L1 loss. Models were trained with CTransPath^20^ and RetCCL^21^ encoders. For a naïve comparator encoder, an Encoder4Editing architecture was trained to minimize an equal ratio of LPIPS and DISTS metrics between the real and generated images from the unmodified StyleGAN2 network, as these yielded the most accurate image representation compared to other non-SSL based comparisons such as L1 loss, L2 loss, and structural similarity. To demonstrate the necessity of the HistoXGAN architecture (which enables direct projection of SSL features into the StyleGAN latent space), we evaluated both an Encoder4Editing and SingleStyle encoder, trained to minimize the same L1 loss from (1). Encoders were trained for 200,000 epochs with a batch size of 8.

### Deep Learning Model Training for Quantitative Assessment

All deep learning models for outcome prediction were trained using the SlideFlow platform. For comparison of predictions of grade, histologic subtype, single gene expression between real / generated images (**Figure 3**, **Supplemental Figure 3**), as well as for illustration of PIK3CA / homologous recombination deficiency (**Figure 4**), and for prediction of tissue source site and ancestry (**Figure 6**), we utilized Xception^47^-based weakly supervised models with ImageNet^8^ pretraining, trained for between 1 to 3 epochs with batch size of 32. For separate visualization of attention and model predictions (**Supplemental Figure 4**), and for prediction of image features for reconstructed histology from MRI (**Figure 6**), we used an attention-MIL architecture trained for 20 epochs. All models were trained with a learning rate of 0.0001 and weight decay of 0.00001. Models for grade, histologic subtype, and single gene expression were trained / evaluated with cross validation for TCGA datasets, and retrained across all of TCGA for application to external datasets; other models were trained across the entire TCGA cohort.

### Visual Representation of Transition Between Histology States

Several approaches are used to demonstrate the robustness of traversing histology feature space within HistoXGAN images. For visualization of grade, histologic subtype, and gene expression patterns (**Figure 3**, **Supplemental Figure 3**), a simple logistic regression model was trained to predict these outcomes using averaged SSL extracted feature vectors across the annotated tumor region from each slide, to obtain a coefficient vector *ν_coef_*. Using a randomly selected baseline feature vector from the corresponding cancer dataset, interpolation is performed between *ν_base_* − *ν_coef_* to *ν_base_* + *ν_coef_*, with images generated at fixed intervals along the interpolation. For visualization along the gradient of a pretrained model *M* (as in **Figure 4** and **Figure 5**), we apply gradient descent to iteratively update a base vector to minimize the loss (2) between the model prediction and the target prediction:

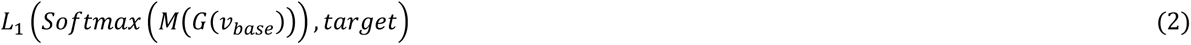

For visualization of principal components of model predictions (**Figure 6**, **Supplemental Figure 6**), a single gradient from the loss as per (2) is generated across a sample of base vectors across the entire dataset. Principal component analysis is used on the resulting gradients to generate 20 orthogonal components *c*_1,2…*n*_ of gradients, and then interpolation is performed between *ν_base_* − *c_i_* to *ν_base_* + *c_i_*.

### Reconstruction of Histology from MRI Images

Dynamic contrast enhanced MRI images acquired on 1.5 T or 3 T magnet strength scanners and digital histology images scanned at 40x with an Aperio AT2 scanner were obtained for 934 patients. Radiomic features were extracted from the region of each DCE-MRI defined by a tumor mask. To generate the tumor masks and visualize results, we employed the previously validated TumorSight Viz platform^48^. Briefly, TumorSight Viz implements a fully automated segmentation approach consisting of a series of convolutional neural networks trained on pre-contrast and post-contrast DCE-MRIs to obtain an initial tumor mask. Following the tumor segmentation process, radiomic features were extracted from the pre-contrast and post-contrast DCE-MRIs along with mathematically computed subtraction and percent enhancement volume maps. Features were also extracted from the peritumoral regions by eroding or dilating the tumor region by approximately 3 mm. Prior to feature extraction, each volume map was transformed into eight additional maps using three-dimensional wavelet filters. Wavelet filters, employing high or low pass filters in each dimension, enhance various frequency components of the volumes and are capable of capturing important textural information^49^. The Pyradiomics library was used to generate the wavelet-transformed volume maps^50^. Once all volume maps were generated, features were extracted from a set of standard feature classes, resulting in a total of 16,379 generated features (**Supplemental Table 12**)^3^.

Histologic features were extracted from tumor tiles from the matched histology samples using an SSL feature extractor, and the average feature vector in the tumor region was calculated for each case. An encoder was trained for 5 epochs with a single leaky ReLU hidden layer to convert MRI radiomic features to SSL histologic features. The MRI encoder was trained with a two-component loss, an L1 loss between the encoder prediction and mean SSL histologic feature vector, and the and an L1 loss between features extracted from an image generated from encoder prediction and the mean SSL feature vector. This was performed across 5 cross folds of the University of Chicago dataset, such that a predicted feature vector was generated for each patient in the dataset. Accuracy of reconstruction was assessed by comparing predictions for grade, tumor histologic subtype, and 775 clinically relevant gene expression features that can be identified from histology for reconstructed images to predictions from the same models applied to the original whole slide images.

Predictive models for these clinically relevant gene expression signatures were trained first using three-fold cross validation in TCGA to verify accuracy of predictions (**Supplemental Figure 6**), with a composite model trained across TCGA for use in assessment of reconstruction from MRI. For comparison, logistic regression models were trained from MRI features using the same cross folds to predict these clinical and gene expression features directly from MRI features.

### Pathologist image interpretation

To assess accuracy of reconstructed images from HistoXGAN and the three comparator encoders, four pathologists were presented with one image from each cancer in the CPTAC validation set, along with reconstructed images from HistoXGAN and alternative encoders presented in a random order. Study pathologists were asked to select the generated image most similar to the original image (i.e. which image would most likely represent a nearby section of the same tumor). This process was repeated with both the CTransPath and RetCCL feature extractors.

To characterize the histologic features identified by deep learning models that were predictive of *PIK3CA* alterations and HRD status, 20 random images were selected from the TCGA-BRCA dataset, and gradient descent was used to alter the base feature vector to produce a high / low likelihood of predicted *PIK3CA* alteration, or high / low HRD score. Images were generated at fixed steps during this transition, and study pathologists were asked to qualitatively describe tumor, cytoplasmic, stromal, immune, and vascular changes that occur during this transition. Features that were consistently identified by the majority of pathologists were selected to identify a consensus for pathologic features representing each of these image transitions.

To verify the veracity of these associations, previously reported annotations^51^ for epithelial, nuclear, and mitotic grade, as well as for necrosis, inflammation, and fibrous foci were compared among cases with or without *PIK*3*CA* alteration or high HRD score. Additionally, to determine the minimum number of annotations that would be needed to confirm these associations with traditional whole slide image review, we repeated these comparisons using 50, 100, 200, 400, or 800 cases of the total TCGA-BRCA cohort, sampled randomly for 100 iterations.

### Statistical analysis

For analysis of the perceptual accuracy of reconstruction of grade, histologic subtype, and single gene expression (**Supplemental Tables 3**, **4, and 5**) – the Pearson correlation coefficient was computed between the averaged model prediction across all tiles from whole slide images, and the averaged prediction across regenerated tiles from extracted features from all tiles across these images. For TCGA cohorts, the predictions were grouped from held out test sets with three-fold cross validation, whereas for CPTAC cohorts the predictions were all made from the same model. For comparison of previously annotated histologic features (such as tubule formation or nuclear pleomorphism) between PIK3CA altered / non-altered and HRD high / low cases, adjusted odds ratios and corresponding Wald statistics were computed for each histologic feature using a multivariable logistic regression for all features. For comparison of predictions from MRI generated images versus real whole slide images, Pearson correlation coefficient was computed, and false discovery rate correction was performed with Benjamini Hochberg^52^ method with false discover / family wide error rate of 5%. Identical analysis was performed for predictions from logistic regression trained from MRI features to directly predict model outputs. All statistical testing performed was two-sided at an *a* = 0.05 level.

## Data Availability

Data from TCGA including digital histology and the majority of clinical annotations used are available from https://portal.gdc.cancer.gov/ and https://cbioportal.org. Annotations for HRD status are available in the published work of Knijnenburg et al^45^, and annotations for genomic ancestry were obtained from Carrot-Zhang et al^53^. Matched MRI features and extracted histology images will be published on acceptance of the manuscript. Source data are provided with this paper.

## Code Availability

Our code used for analysis from this project is available from https://github.com/fmhoward/HistoXGAN.

## Author Contributions

F.M.H. was responsible for concept proposal, study design, and essential programming work. K.C., M.S., J.V., and G.K. performed blinded assessment of similarity of generated / real pathologic images, and interpretation of relevant features from generated histology. C.F. and C.M.P. computed biologically relevant gene expression signatures in the TCGA dataset. All authors contributed to the data analysis and writing of the manuscript.

## Acknowledgments

F.M.H. received support from the NIH/NCI (K08CA283261) and the Cancer Research Foundation. A.T.P received support from the NIH/NIDCR (R56DE030958), the European Comission Horizon (2021-SC1-BHC), NIH/NCI (R01CA276652), SU2C, the Adenoid Cystic Carcinoma Research Foundation, the Cancer Research Foundation, and the American Cancer Society. A.T.P., D.H., and F.M.H. received support from the Department of Defense (BC211095P1). D.H. received support from the NIH/NCI (1P20-CA233307) and the Breast Cancer Research Foundation (BCRF-21-071). C.M.P received support from the NCI Breast SPORE program (P50-CA058223), and the Breast Cancer Research Foundation (BCRF-23-127).

---

**Supplemental Figure 1.**
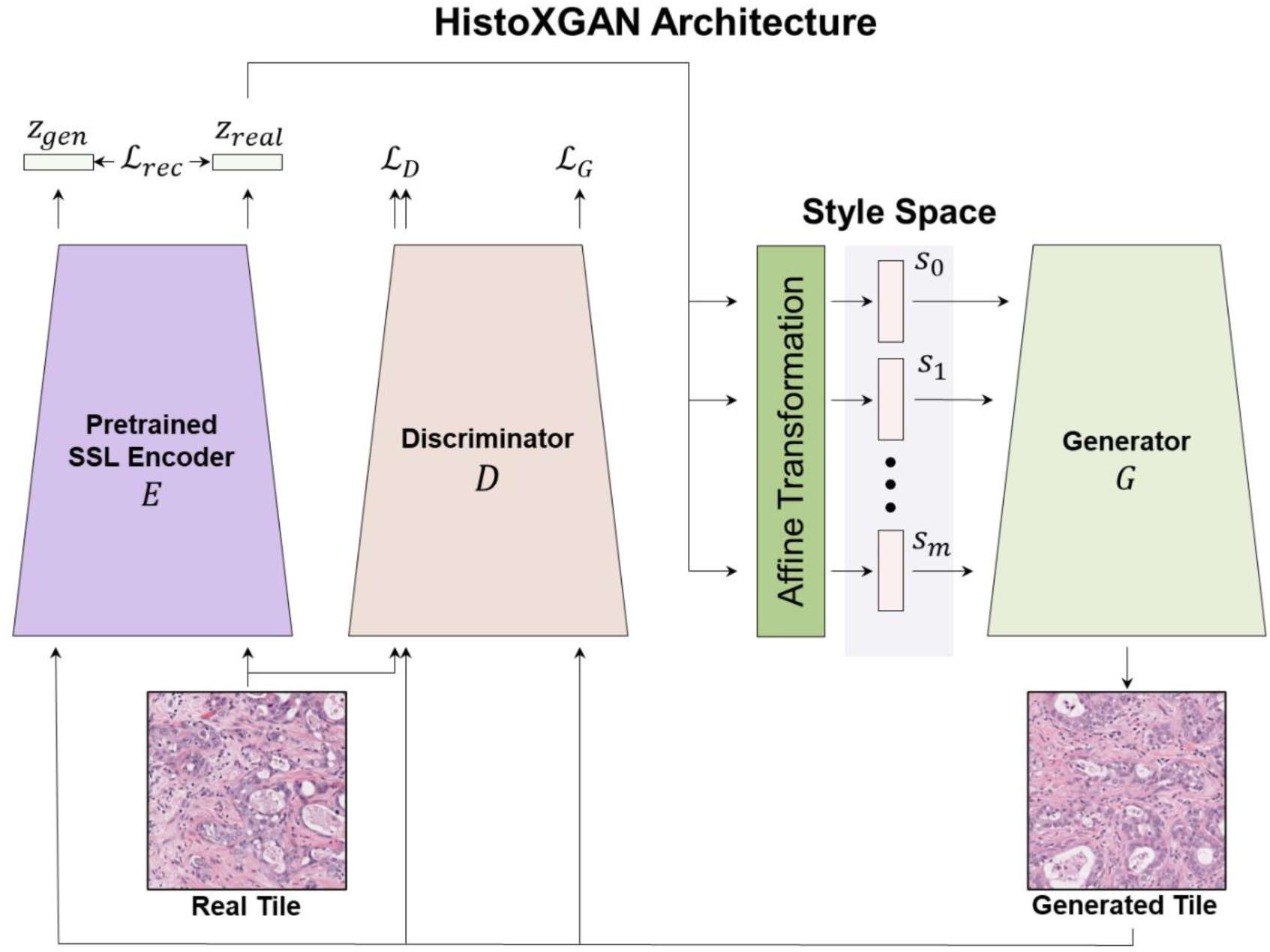
Overview of Generator Architecture and Study Analysis. The HistoXGAN architecture is based on StyleGAN2, with simultaneous training of a generator *G* and discriminator *D*. During training, latent image features (z) are extracted using a pretrained self-supervised feature encoder *E*. These latent features are transformed into style vectors (*s_0_*, …, *s*_n_) which are used to recreate a pathology image tile. An additional L1 loss ℒ*_rec_* is added for the comparison of the encoded features from the reconstructed pathology tile *z_gen_* and the input tile *z_real_*.

**Supplemental Figure 2.**
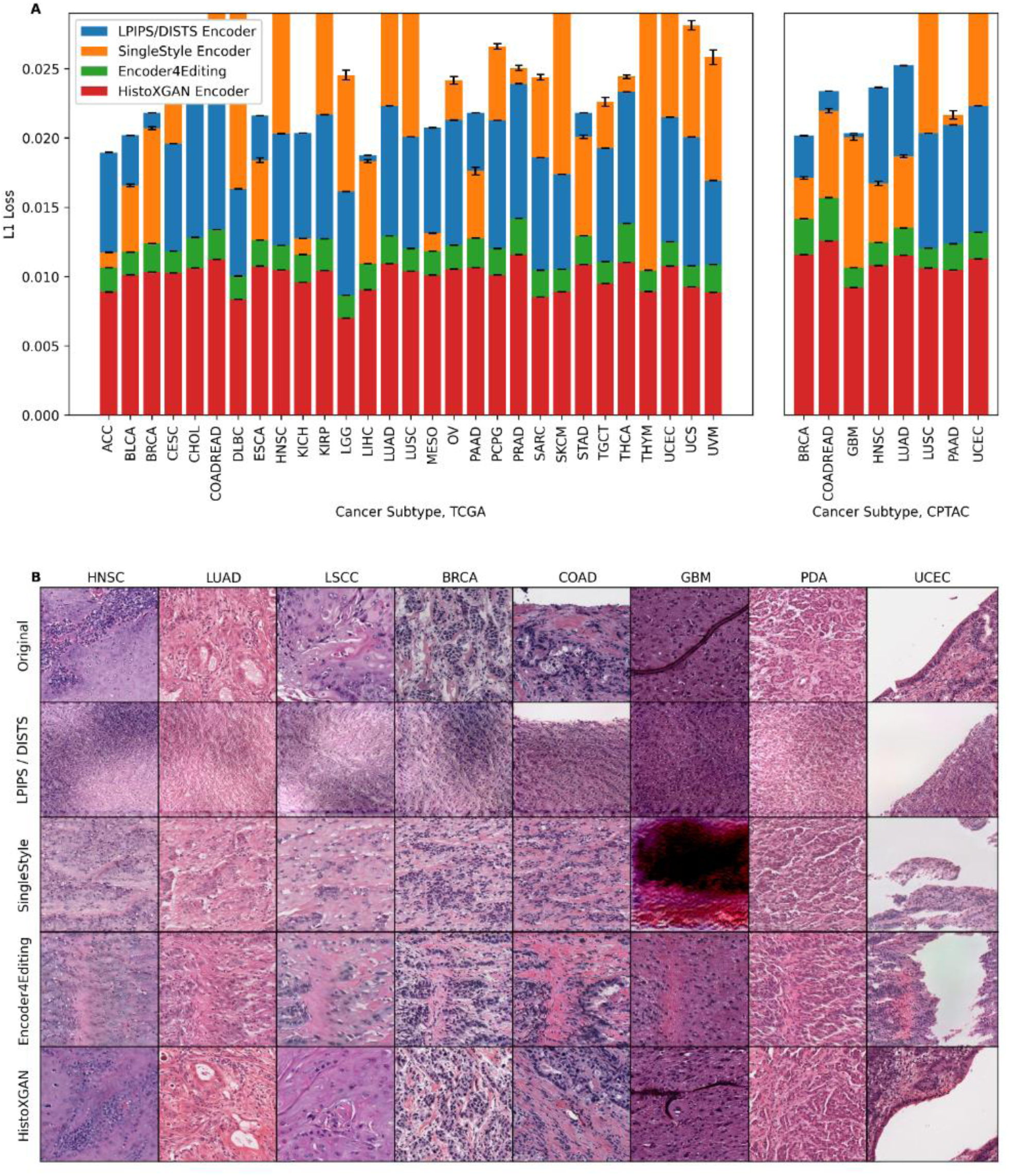
Reconstruction Accuracy in Training and Validation Datasets for RetCCL Encoders. We compare reconstruction accuracy from the real and reconstructed images for HistoXGAN and other architectures for embedding images in GAN latent space. For comparison, we use encoders designed to recreate images from a StyleGAN2 model trained identically to the HistoXGAN model. The Learned Perceptual Image Patch Similarity (LPIPS) / Deep Image Structure and Texture Similarity (DISTS) encoder uses an equal ratio of LPIPS / DISTS loss between the real and reconstructed images to train the encoder. The Single Layer and Encoder4Editing encoders are trained to minimize L1 loss between RetCCL feature vector of the real and reconstructed images. **A.** HistoXGAN provides more accurate reconstruction of RetCCL features across the TCGA dataset used for GAN training (n = 8,120) and CPTAC validation (n = 1,328) dataset, achieving an average of 30% improvement in L1 loss over the Encoder4Editing encodings in the validation dataset. **B.** HistoXGAN reconstructed images consistently provided more accurate representations of features from the input image across cancer types in the CPTAC validation dataset.

**Supplemental Figure 3.**
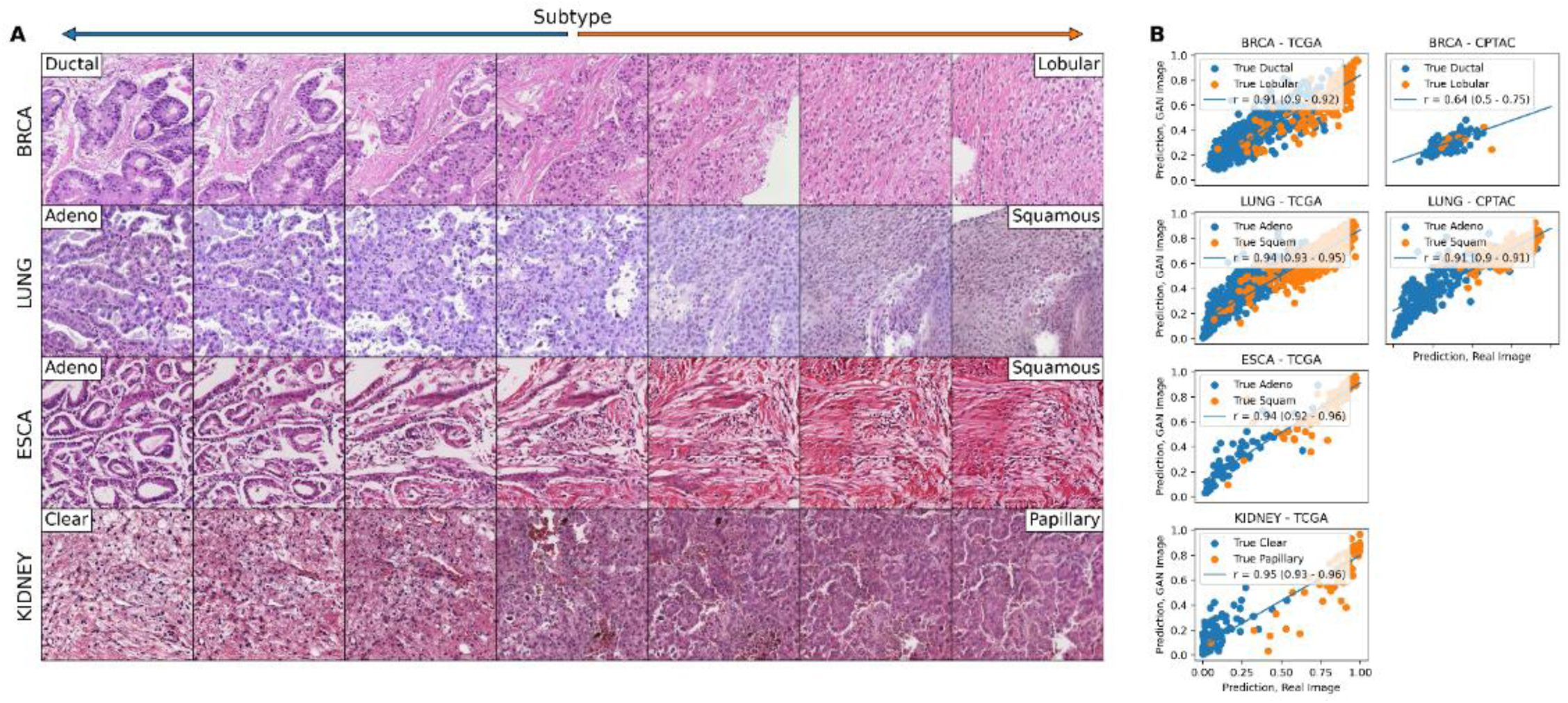
Perceptual Consistency of Histologic Subtype in Reconstructed Images Across Cancer Types. **A.** Illustration of transition between tumor histologic subtypes across a single image from four cancer types. A vector differentiating the two most frequent subtype classes is derived from the coefficients of a logistic regression predicting subtype from CTransPath features. This vector is subtracted from the base image to visualize the first class, and added to the base image to visualize the second class. **B.** Correlation between predictions of histologic subtype from real and reconstructed tiles, averaged per patient, across cancer types, demonstrating a high perceptual similarity of the histologic subtype of the real and generated images. For the TCGA datasets, a deep learning model was trained to predict subtype from real tiles for each cancer type using three-fold cross validation. The correlation between predictions for real / generated images is aggregated for the three held out validation sets. For the CPTAC validation, a deep learning model trained across the entire corresponding TCGA dataset was used to generate predictions. True pathologist determined subtype is annotated on the images when available.

**Supplemental Figure 4.**
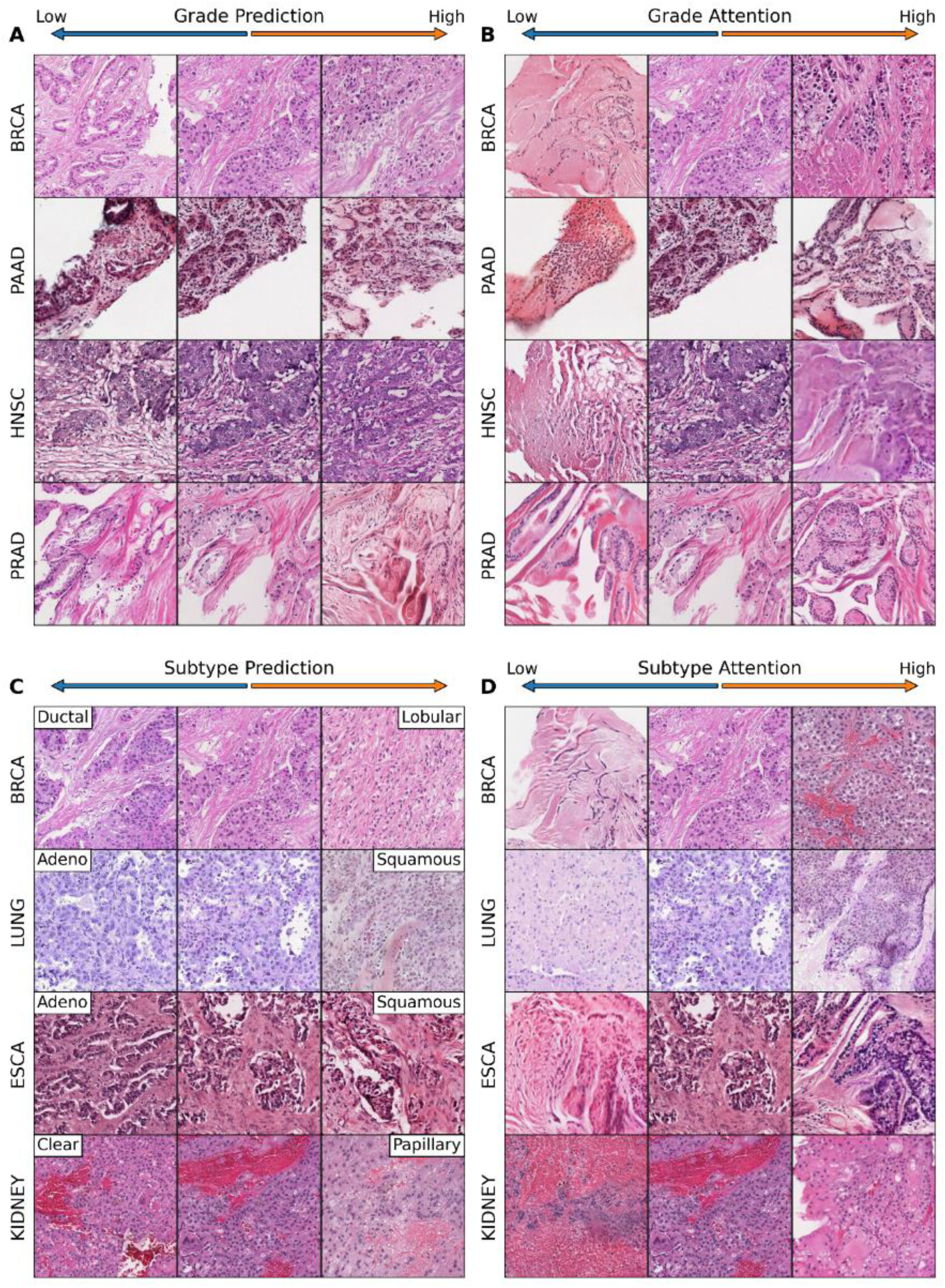
Interpretability of Attention-based / Multiple Instance Learning Models. We demonstrate that this HistoXGAN approach can be used to understand attention-based multiple instance learning (MIL) approaches or other multi-step models. Attention-MIL models were trained to predict grade and histologic subtype across multiple cancers in TCGA. Image tiles were generated with HistoXGAN, and gradient descent was used to perturb the image features to increase the predicted likelihood of grade / subtype class or increase attention towards the image tile. Grade and subtype prediction demonstrated expected histologic findings, and tiles demonstrated a more fibrous and acellular appearance as attention decreased.

**Supplemental Figure 5.**
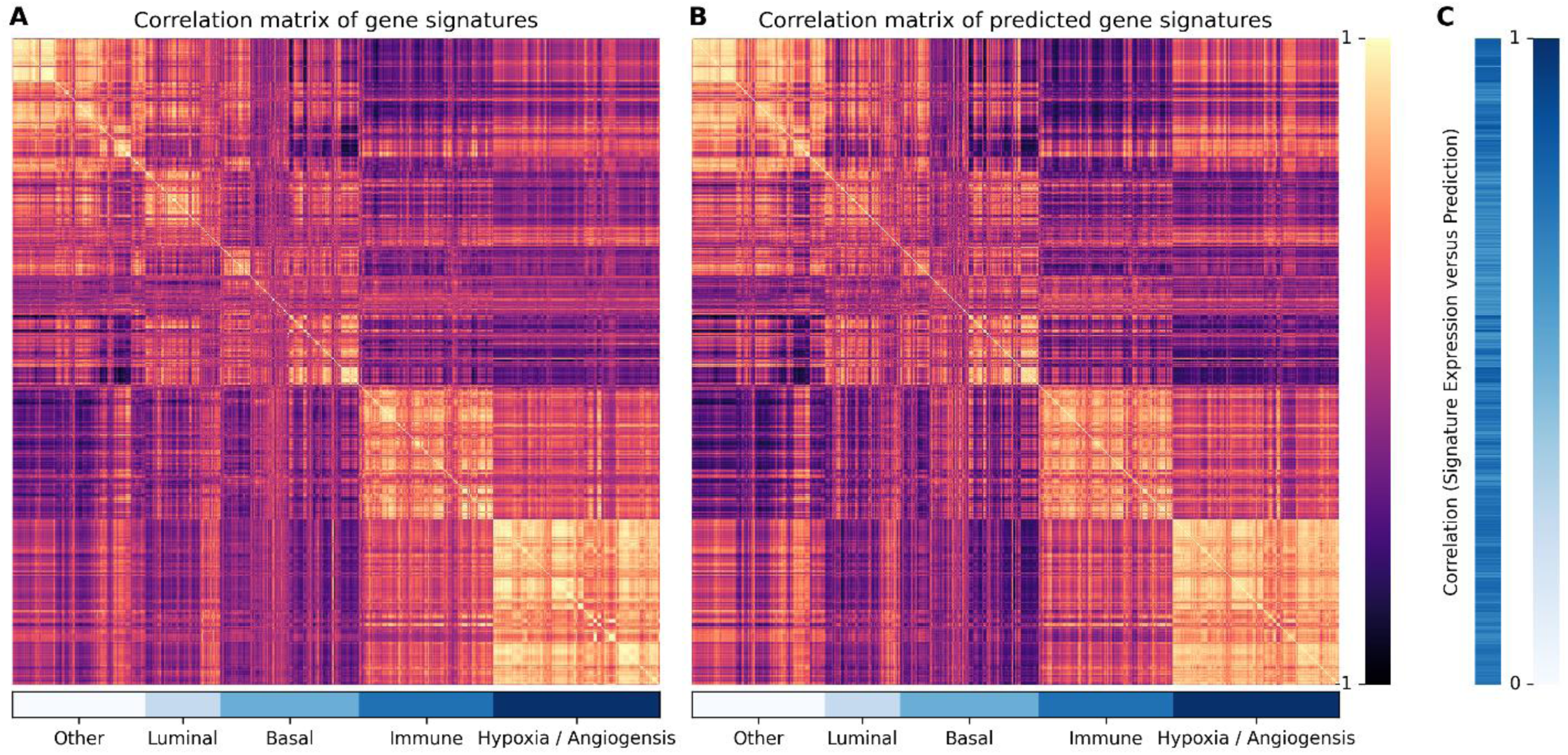
Validating Predicted Gene Signatures as a Method of Assessing the Biologic Similarity of Histologic Images. Deep learning models were trained with three-fold cross validation to predict 775 clinically relevant gene expression signatures from digital histology in the TCGA-BRCA cohort, and predictions in the held-out test set were compared to the true values of gene expression from RNA sequencing. **A.** Correlation matrix (illustrating Pearson correlation coefficient) of gene signatures in the TCGA cohort, demonstrating that the 775 signatures yield several orthogonal groups of highly correlated signatures, illustrating some degree of redundancy – especially in the immune / hypoxia signatures. **B.** Correlation matrix of the gene signatures predicted from digital histology in the pooled held-out cohorts, demonstrating a highly similar pattern of signature grouping to the true gene expression signatures. **C.** Pearson correlation between the gene signatures calculated from RNA sequencing, and the predicted signatures in the pooled held-out cohorts. Clinically relevant gene expression signatures could be accurately predicted from histology, with an average correlation coefficient between real and predicted signature of 0.45.

**Supplemental Figure 6.**
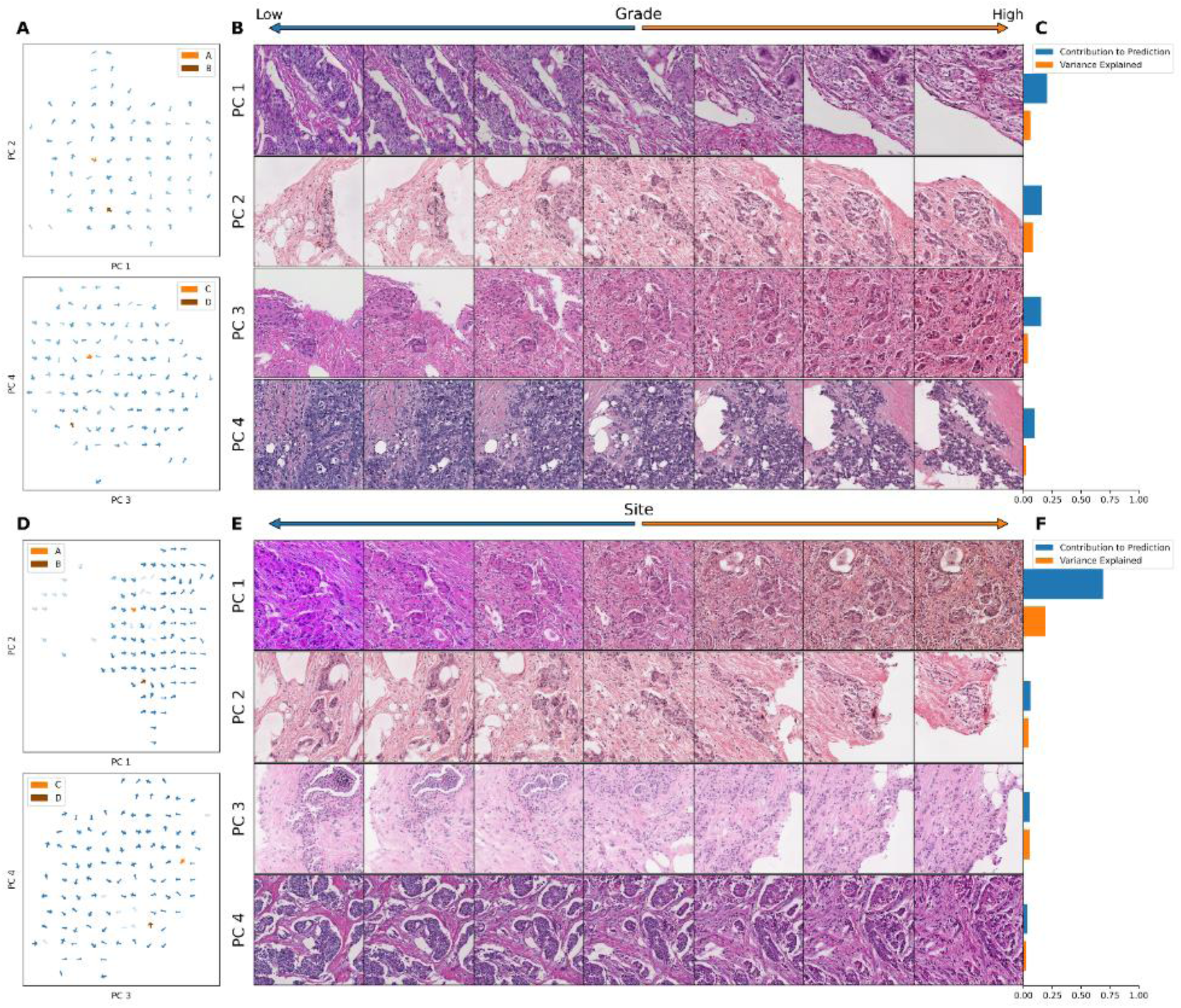
Identifying Orthogonal Features that Contribute to Model Predictions. Tile based weakly supervised models were trained to predict grade and contributing site for TCGA-BRCA (n = 943). The gradient with respect to a prediction of these outcomes (grade low / high; site A / B) was calculated for the average feature vector across each slide in the dataset. Principal component analysis was applied to these gradients, and components were sorted by the magnitude of difference of the component between gradients toward each outcome class, i.e. the strength of the contribution of the component to the prediction. **A, D.** Gradients to predict high grade / site B were projected into spaces of pairs of principal components. Magnitude of the gradient is indicated by arrow thickness. Whereas the component that led to a stronger prediction of high / low grade was highly variable, for nearly all images application of the first principal component led to a stronger prediction for site B (i.e. all arrows pointing to the right in the first plot in figure D). The tile with the maximum gradient for each component is selected for plotting (labeled A - D for principle components 1 – 4). **B, E.** The top four principles components for grade and site were used to perturb images. The first principal component in site clearly caused a shift in the stain color of the image. **C, F.** The relative contribution of the component to the gradient and the proportion of variance explained by the component are listed. Approximately 70% of the gradient towards a higher prediction could be explained by the first principal component for site, illustrating the site model is relatively simple and based nearly entirely on stain color difference.

**Supplemental Table 1.** Reconstruction Accuracy in Training and Validation Datasets for CTransPath Encoders. We compare reconstruction accuracy from the real and reconstructed images for HistoXGAN and other architectures for embedding images in GAN latent space. For comparison, we use encoders designed to recreate images from a StyleGAN2 model trained identically to the HistoXGAN model. The Learned Perceptual Image Patch Similarity (LPIPS) / Deep Image Structure and Texture Similarity (DISTS) encoder uses an equal ratio of LPIPS / DISTS loss between the real and reconstructed images to train the encoder. The Single Layer and Encoder4Editing encoders are trained to minimize L1 loss between CTransPath feature vector of the real and reconstructed images.

| Source |  | n | n Tiles | LPIP / DISTS<br>L1 Loss | Single Layer<br>L1 Loss | Encoder4Editing<br>L1 Loss | HistoXGAN<br>L1 Loss |
| --- | --- | --- | --- | --- | --- | --- | --- |
| TCGA | ACC | 56 | 78789 | 0.086 (0.010) | 0.054 (0.010) | 0.045 (0.006) | 0.035 (0.004) |
| TCGA | BLCA | 378 | 342463 | 0.082 (0.009) | 0.054 (0.015) | 0.044 (0.006) | 0.034 (0.004) |
| TCGA | BRCA | 943 | 454985 | 0.087 (0.010) | 0.055 (0.017) | 0.044 (0.006) | 0.033 (0.004) |
| TCGA | CESC | 267 | 165088 | 0.083 (0.010) | 0.063 (0.021) | 0.044 (0.006) | 0.034 (0.004) |
| TCGA | CHOL | 38 | 51414 | 0.087 (0.009) | 0.057 (0.019) | 0.044 (0.006) | 0.034 (0.004) |
| TCGA | COADREAD | 428 | 195493 | 0.092 (0.013) | 0.071 (0.023) | 0.046 (0.007) | 0.035 (0.004) |
| TCGA | DLBC | 43 | 32073 | 0.076 (0.011) | 0.063 (0.022) | 0.045 (0.010) | 0.032 (0.005) |
| TCGA | ESCA | 147 | 99625 | 0.085 (0.010) | 0.057 (0.017) | 0.045 (0.007) | 0.035 (0.005) |
| TCGA | HNSC | 401 | 166061 | 0.083 (0.009) | 0.060 (0.019) | 0.045 (0.006) | 0.035 (0.004) |
| TCGA | KICH | 101 | 105461 | 0.094 (0.010) | 0.056 (0.009) | 0.048 (0.006) | 0.036 (0.005) |
| TCGA | KIRP | 270 | 234740 | 0.091 (0.013) | 0.070 (0.022) | 0.049 (0.007) | 0.036 (0.005) |
| TCGA | LGG | 464 | 155579 | 0.081 (0.009) | 0.053 (0.014) | 0.044 (0.006) | 0.031 (0.005) |
| TCGA | LIHC | 359 | 331769 | 0.090 (0.013) | 0.058 (0.019) | 0.045 (0.006) | 0.033 (0.004) |
| TCGA | LUAD | 467 | 335499 | 0.088 (0.010) | 0.062 (0.019) | 0.046 (0.007) | 0.035 (0.004) |
| TCGA | LUSC | 474 | 370542 | 0.083 (0.009) | 0.061 (0.019) | 0.045 (0.006) | 0.035 (0.004) |
| TCGA | MESO | 73 | 42242 | 0.083 (0.009) | 0.053 (0.011) | 0.044 (0.006) | 0.034 (0.004) |
| TCGA | OV | 104 | 108620 | 0.085 (0.010) | 0.061 (0.019) | 0.045 (0.007) | 0.035 (0.005) |
| TCGA | PAAD | 168 | 119144 | 0.085 (0.009) | 0.053 (0.012) | 0.046 (0.007) | 0.035 (0.005) |
| TCGA | PCPG | 173 | 207803 | 0.083 (0.009) | 0.066 (0.022) | 0.044 (0.006) | 0.035 (0.004) |
| TCGA | PRAD | 394 | 202439 | 0.091 (0.010) | 0.057 (0.015) | 0.046 (0.007) | 0.034 (0.004) |
| TCGA | SARC | 250 | 326262 | 0.084 (0.011) | 0.059 (0.019) | 0.047 (0.007) | 0.033 (0.005) |
| TCGA | SKCM | 418 | 322527 | 0.080 (0.009) | 0.063 (0.023) | 0.044 (0.006) | 0.034 (0.004) |
| TCGA | STAD | 371 | 283734 | 0.086 (0.011) | 0.055 (0.014) | 0.046 (0.008) | 0.035 (0.005) |
| TCGA | TGCT | 129 | 109128 | 0.082 (0.008) | 0.057 (0.015) | 0.045 (0.007) | 0.034 (0.005) |
| TCGA | THCA | 480 | 279362 | 0.091 (0.010) | 0.065 (0.022) | 0.047 (0.007) | 0.035 (0.005) |
| TCGA | THYM | 114 | 157626 | 0.078 (0.010) | 0.080 (0.028) | 0.045 (0.008) | 0.033 (0.006) |
| TCGA | UCEC | 477 | 351784 | 0.084 (0.010) | 0.071 (0.025) | 0.045 (0.006) | 0.035 (0.004) |
| TCGA | UCS | 53 | 67853 | 0.082 (0.009) | 0.061 (0.019) | 0.045 (0.006) | 0.035 (0.005) |
| TCGA | UVM | 80 | 35766 | 0.075 (0.009) | 0.063 (0.020) | 0.045 (0.007) | 0.033 (0.005) |
| CPTAC | BRCA | 105 | 56318 | 0.084 (0.008) | 0.067 (0.016) | 0.053 (0.008) | 0.040 (0.005) |
| CPTAC | COADREAD | 104 | 76915 | 0.086 (0.009) | 0.071 (0.015) | 0.056 (0.007) | 0.041 (0.005) |
| CPTAC | GBM | 177 | 143858 | 0.080 (0.010) | 0.062 (0.016) | 0.048 (0.007) | 0.038 (0.006) |
| CPTAC | HNSC | 108 | 35064 | 0.080 (0.009) | 0.062 (0.018) | 0.049 (0.007) | 0.038 (0.005) |
| CPTAC | LUAD | 221 | 425473 | 0.087 (0.009) | 0.057 (0.012) | 0.048 (0.006) | 0.038 (0.004) |
| CPTAC | LUSC | 202 | 359394 | 0.081 (0.008) | 0.066 (0.021) | 0.046 (0.006) | 0.036 (0.004) |
| CPTAC | PAAD | 164 | 99093 | 0.082 (0.009) | 0.059 (0.016) | 0.049 (0.008) | 0.037 (0.006) |
| CPTAC | UCEC | 247 | 184767 | 0.083 (0.010) | 0.074 (0.021) | 0.049 (0.008) | 0.038 (0.006) |

**Supplemental Table 2.**
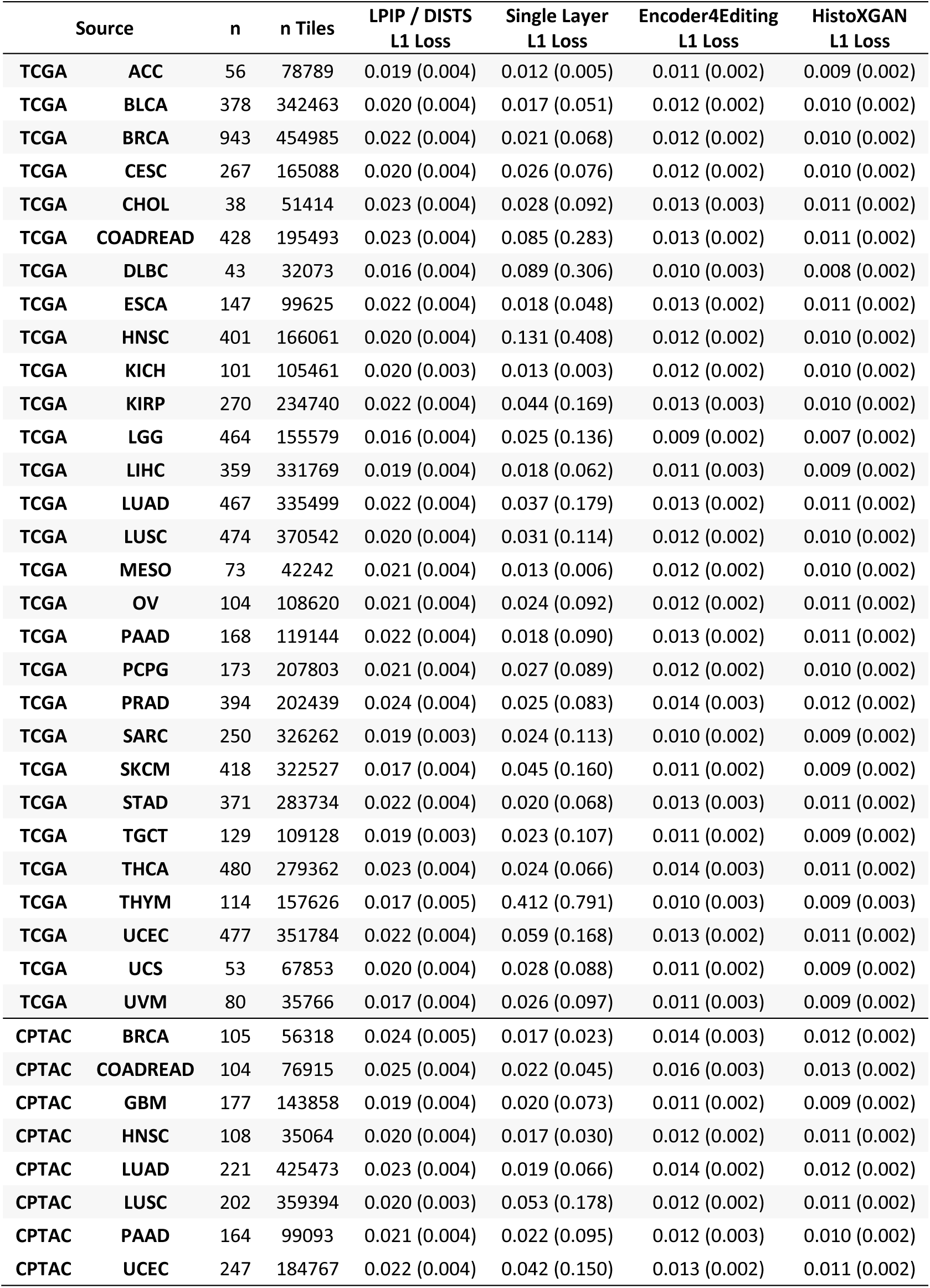
Reconstruction Accuracy in Training and Validation Datasets for RetCCL Encoders. We compare reconstruction accuracy from the real and reconstructed images for HistoXGAN and other architectures for embedding images in GAN latent space. For comparison, we use encoders designed to recreate images from a StyleGAN2 model trained identically to the HistoXGAN model. The Learned Perceptual Image Patch Similarity (LPIPS) / Deep Image Structure and Texture Similarity (DISTS) encoder uses an equal ratio of LPIPS / DISTS loss between the real and reconstructed images to train the encoder. The Single Layer and Encoder4Editing encoders are trained to minimize L1 loss between RetCCL feature vector of the real and reconstructed images.

**Supplemental Table 3:**
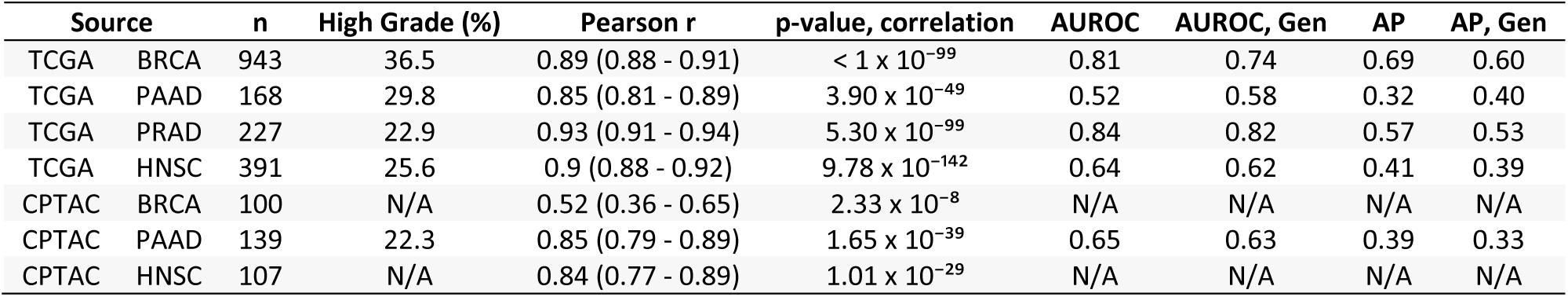
Perceptual Consistency of Tumor Grade in Reconstructed Images Across Cancer Types. Correlation between predictions of grade from real and reconstructed tiles, averaged per patient, across cancer types, demonstrating a high perceptual similarity of the grade of the real and generated images. For the TCGA datasets, a deep learning model was trained to predict grade from real tiles for each cancer type using three-fold cross validation. The correlation between predictions for real / generated images is aggregated for the three held out validation sets. For the CPTAC validation, a deep learning model trained across the entire corresponding TCGA dataset was used to generate predictions. Average area under the receiver operating characteristic (AUROC) and average precision (AP) are listed for prediction of grade using the above models, as well as when predictions from these models are made with reconstructed (Gen) versions of tiles. The similar AUROC / AP from real tiles and reconstructed tiles illustrates the reconstructed tiles retain informative data with regards to grade.

| Source |  | n | High Grade (%) | Pearson r | p-value, correlation | AUROC | AUROC, Gen | AP | AP, Gen |
| --- | --- | --- | --- | --- | --- | --- | --- | --- | --- |
| TCGA | BRCA | 943 | 36.5 | 0.89 (0.88 - 0.91) | $< 1 \times 10^{-99}$ | 0.81 | 0.74 | 0.69 | 0.60 |
| TCGA | PAAD | 168 | 29.8 | 0.85 (0.81 - 0.89) | $3.90 \times 10^{-49}$ | 0.52 | 0.58 | 0.32 | 0.40 |
| TCGA | PRAD | 227 | 22.9 | 0.93 (0.91 - 0.94) | $5.30 \times 10^{-99}$ | 0.84 | 0.82 | 0.57 | 0.53 |
| TCGA | HNSC | 391 | 25.6 | 0.9 (0.88 - 0.92) | $9.78 \times 10^{-142}$ | 0.64 | 0.62 | 0.41 | 0.39 |
| CPTAC | BRCA | 100 | N/A | 0.52 (0.36 - 0.65) | $2.33 \times 10^{-8}$ | N/A | N/A | N/A | N/A |
| CPTAC | PAAD | 139 | 22.3 | 0.85 (0.79 - 0.89) | $1.65 \times 10^{-39}$ | 0.65 | 0.63 | 0.39 | 0.33 |
| CPTAC | HNSC | 107 | N/A | 0.84 (0.77 - 0.89) | $1.01 \times 10^{-29}$ | N/A | N/A | N/A | N/A |

**Supplemental Table 4:**
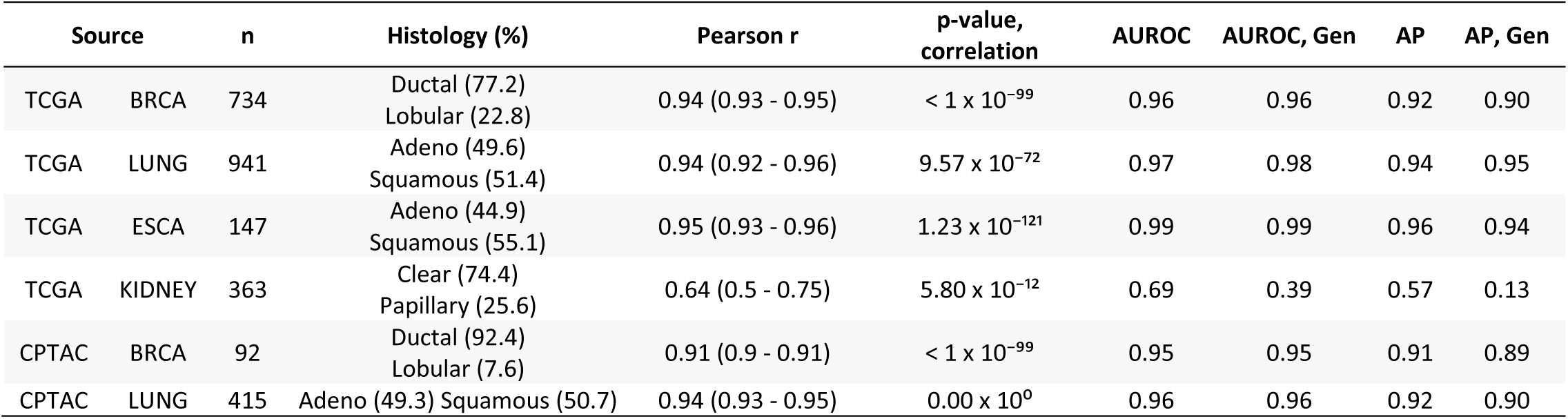
Perceptual Consistency of Histologic Subtype in Reconstructed Images Across Cancer Types. Correlation between predictions of histologic subtype from real and reconstructed tiles, averaged per patient, across cancer types, demonstrating a high perceptual similarity of the histologic subtype of the real and generated images. For the TCGA datasets, a deep learning model was trained to predict subtype from real tiles for each cancer type using three-fold cross validation. The correlation between predictions for real / generated images is aggregated for the three held out validation sets. For the CPTAC validation, a deep learning model trained across the entire corresponding TCGA dataset was used to generate predictions. Average area under the receiver operating characteristic (AUROC) and average precision (AP) are listed for prediction of histologic subtype using the above models, as well as when predictions from these models are made with reconstructed (Gen) versions of tiles. The similar AUROC / AP from real tiles and reconstructed tiles illustrates the reconstructed tiles retain informative data with regards to histologic subtype.

**Supplemental Table 5:** Perceptual Consistency of Gene Expression in Reconstructed Images. Correlation between predictions of gene expression from real and reconstructed tiles, averaged per patient, demonstrating a high perceptual similarity of the gene expression of the real and generated images. For TCGA, a deep learning model was trained to predict gene expression from real tiles from TCGA-BRCA using three-fold cross validation. The correlation between predictions for real / generated images is aggregated for the three held out validation sets. For the CPTAC-BRCA validation, a deep learning model trained across the entire TCGA-BRCA dataset was used to generate predictions. Also listed are the correlation between model predictions and true gene expression, as well as the same correlations made from the reconstructed (Gen) tiles. The similar correlation coefficients from real tiles and reconstructed tiles illustrates the reconstructed tiles retain informative data with regards to histologic subtype.

| Source | Gene | n | Pearson r, Real vs Gen | p-value | Pearson r, Real vs Expression | p-value | Pearson r, Gen vs Expression | p-value |
| --- | --- | --- | --- | --- | --- | --- | --- | --- |
| TCGA | CD3G | 941 | 0.9 (0.88 - 0.91) | $< 1 \times 10^{-100}$ | 0.45 (0.39 - 0.49) | $< 1 \times 10^{-100}$ | 0.38 (0.32 - 0.43) | $< 1 \times 10^{-100}$ |
| TCGA | COL1A1 | 941 | 0.84 (0.82 - 0.86) | $6.88 \times 10^{-269}$ | 0.45 (0.4 - 0.5) | $< 1 \times 10^{-100}$ | 0.31 (0.26 - 0.37) | $< 1 \times 10^{-100}$ |
| TCGA | MKI67 | 941 | 0.91 (0.9 - 0.92) | $0.00 \times 10^0$ | 0.43 (0.38 - 0.48) | $< 1 \times 10^{-100}$ | 0.35 (0.3 - 0.41) | $< 1 \times 10^{-100}$ |
| TCGA | EPCAM | 941 | 0.9 (0.88 - 0.91) | $0.00 \times 10^0$ | 0.32 (0.26 - 0.37) | $< 1 \times 10^{-100}$ | 0.25 (0.19 - 0.31) | $< 1 \times 10^{-100}$ |
| CPTAC | CD3G | 97 | 0.73 (0.62 - 0.81) | $3.08 \times 10^{-17}$ | 0.3 (0.1 - 0.47) | $< 1 \times 10^{-100}$ | 0.21 (0.02 - 0.4) | 0.04 |
| CPTAC | COL1A1 | 97 | 0.74 (0.64 - 0.82) | $2.89 \times 10^{-18}$ | 0.56 (0.4 - 0.68) | $< 1 \times 10^{-100}$ | 0.55 (0.39 - 0.67) | $< 1 \times 10^{-100}$ |
| CPTAC | MKI67 | 97 | 0.82 (0.75 - 0.88) | $5.34 \times 10^{-25}$ | 0.04 (-0.16 - 0.24) | 0.67 | 0.07 (-0.13 - 0.26) | 0.51 |
| CPTAC | EPCAM | 97 | 0.44 (0.27 - 0.59) | $5.21 \times 10^{-6}$ | 0.1 (-0.1 - 0.29) | 0.33 | -0.04 (-0.24 - 0.16) | 0.69 |

**Supplemental Table 6:** Correlation of Annotated Histologic Features with *PIK3CA* mutational status. Previously reported annotations for epithelial, nuclear, and mitotic grade, as well as for necrosis, fibrous foci, and inflammation were compared between cases with / without *PIK3CA* mutations. Adjusted odds ratio was computed for the association of each feature with *PIK3CA* (association listed for high tubule formation score, high nuclear pleomorphism score, high mitosis score, and present necrosis, fibrosis, and inflammation). Additionally, to determine how many annotated cases would be needed to identify these correlations, we calculated these adjusted odds ratios for subsets of 50, 100, 200, and 400, 800 cases of the whole cohort, with these subsets selected at random and repeated 100-fold. The median odds ratio over these 100 iterations is listed below.

| Feature | Adjusted Odds Ratio (95% CI) | z-stat | p-value | Median adjusted odds ratio (95% CI) computed with 100 iterations from subset of cases |  |  |  |  |
| --- | --- | --- | --- | --- | --- | --- | --- | --- |
|  |  |  |  | 50 cases | 100 cases | 200 cases | 400 cases | 800 cases |
| Decreased Tubule Formation | 0.6 (0.44 - 0.82) | -3.22 | 0.001 | 0.61 (0.16 - 2.28) | 0.64 (0.24 - 1.67) | 0.62 (0.30 - 1.27) | 0.61 (0.38 - 1.00) | 0.60 (0.43 - 0.84) |
| Nuclear Pleomorphism | 0.61 (0.43 - 0.87) | -2.77 | 0.006 | 0.55 (0.07 - 4.48) | 0.61 (0.20 - 1.83) | 0.59 (0.27 - 1.25) | 0.60 (0.35 - 1.03) | 0.62 (0.42 - 0.90) |
| Mitosis | 0.86 (0.58 - 1.28) | -0.75 | 0.45 | 1.0 (0.14 - 7.35) | 0.82 (0.23 - 2.99) | 0.90 (0.38 - 2.12) | 0.88 (0.48 - 1.62) | 0.86 (0.56 - 1.32) |
| Necrosis | 0.58 (0.4 - 0.84) | -2.9 | 0.004 | 0.57 (0.08 - 4.27) | 0.57 (0.17 - 1.89) | 0.59 (0.26 - 1.34) | 0.59 (0.32 - 1.09) | 0.59 (0.39 - 0.88) |
| Fibrous Focus | 1.08 (0.79 - 1.47) | 0.49 | 0.62 | 1.12 (0.23 - 5.52) | 1.06 (0.35 - 3.26) | 1.04 (0.48 - 2.25) | 1.07 (0.66 - 1.73) | 1.09 (0.77 - 1.53) |
| Inflammation | 1.02 (0.73 - 1.43) | 0.1 | 0.92 | 1.15 (0.21 - 6.46) | 1.08 (0.35 - 3.36) | 1.03 (0.50 - 2.14) | 0.98 (0.58 - 1.66) | 1.01 (0.69 - 1.48) |

**Supplemental Table 7:**
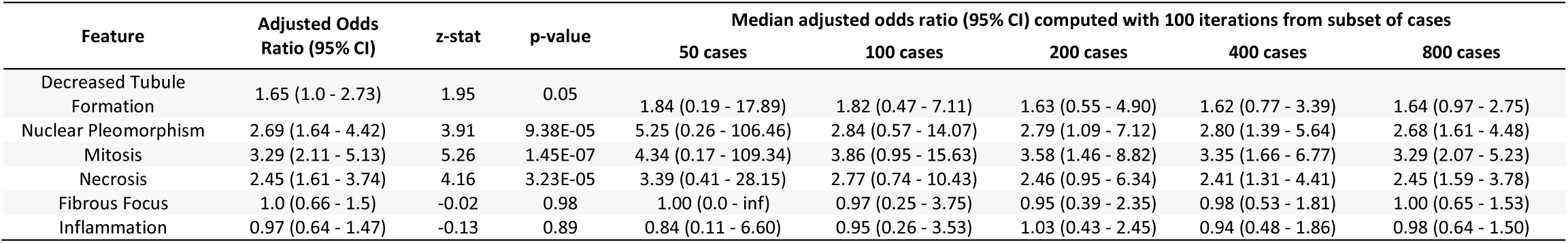
Correlation of Annotated Histologic Features with Homologous Recombination Deficiency. Previously reported annotations for epithelial, nuclear, and mitotic grade, as well as for necrosis, fibrous foci, and inflammation were compared between cases with high / low HRD scores. Adjusted odds ratio was computed for the association of each feature with HRD status (association listed for high tubule formation score, high nuclear pleomorphism score, high mitosis score, and present necrosis, fibrosis, and inflammation). Additionally, to determine how many annotated cases would be needed to identify these correlations, we calculated these adjusted odds ratios for subsets of 50, 100, 200, and 400, 800 cases of the whole cohort, with these subsets selected at random and repeated 100-fold. The median odds ratio over these 100 iterations is listed below.

**Supplemental Table 10:** Homogenous Loss for Models Predicting Tissue Source Site. Tile based weakly supervised models were trained to predict site four datasets within TCGA. The gradient with respect to site prediction was calculated for the average feature vector across each slide in the dataset. Principal component analysis was applied to these gradients, and components were sorted both by the magnitude of difference of the component between gradients toward each outcome class, i.e. the strength of the contribution of the component to the prediction. This process was repeated when applying Reinhard and CycleGAN normalization. Predictions are dominated by a single uniform principal component without normalization and with Reinhard normalization, but this effect is largely mitigated by CycleGAN normalization.

| Source | n | Without Normalization |  | Reinhard Normalized |  | CycleGAN Normalized |  |
| --- | --- | --- | --- | --- | --- | --- | --- |
|  |  | Contribution (Std) | Variance Explained | Contribution (Std) | Variance Explained | Contribution (Std) | Variance Explained |
| BRCA | 943 | 0.691 (0.937) | 0.189 | 0.616 (0.562) | 0.28 | 0.160 (0.830) | 0.068 |
| BLCA | 378 | 0.686 (0.711) | 0.28 | 0.402 (1.711) | 0.381 | 0.258 (1.586) | 0.277 |
| COADREAD | 428 | 0.566 (0.713) | 0.166 | 0.315 (0.636) | 0.098 | 0.186 (0.402) | 0.052 |
| ESCA | 147 | 0.207 (0.323) | 0.067 | 0.164 (0.637) | 0.169 | 0.442 (0.580) | 0.118 |

**Supplemental Table 11:**
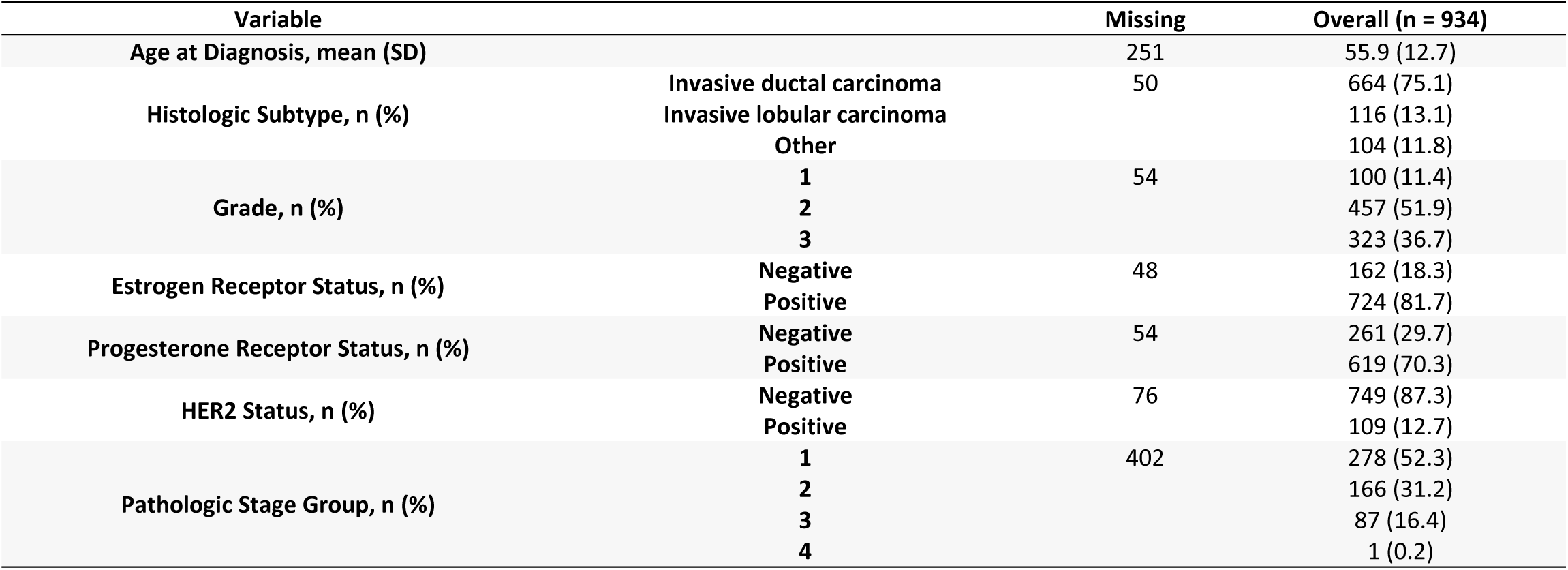
Demographics of Included Patients from University of Chicago with Dynamic Contrast Enhanced Magnetic Resonance Imaging and Digital Histology.

**Supplemental Table 12:** Feature Classes Extracted from MRI Images.

| Class | Number of Features | Description |
| --- | --- | --- |
| Shape-Based | 14 | Describe the size and shape of the region of interest using the segmentation mask and do not depend on gray-scale values. Examples include sphericity and compactness. |
| First Order Statistics | 3,348 | Describe the distribution of voxel intensities within the region of interest. Examples include mean, minimum, and maximum gray-scale values. |
| Gray Level Co-occurrence Matrix | 4,462 | Describe the distribution of co-occurrence of gray-scale values at a given offset distance. Examples include joint average and correlation. |
| Gray Level Run Length Matrix | 2,960 | Quantify runs of consecutive pixels with the same gray level value. Examples include short run emphasis and long run emphasis. |
| Gray Level Size Zone Matrix | 2,960 | Describe the number of zones of voxels with the same gray-scale values. Examples include small area emphasis and large area emphasis. |
| Neighboring Gray Tone Difference Matrix | 45 | Describe the difference between the gray-scale values of a voxel and its neighbors up to a specified distance. Examples include coarseness and busyness. |
| Gray Level Dependence Matrix | 2,590 | Describe the number of voxels that are dependent on the center voxel. Two voxels are dependent if their gray-scale value differences are below a specified threshold. |

